# Allosteric and Energetic Remodeling by Protein Domain Extensions

**DOI:** 10.1101/2025.07.24.666590

**Authors:** Cristina Hidalgo-Carcedo, Andre J. Faure, Aina Martí-Aranda, Taraneh Zarin, Ben Lehner

**Affiliations:** Centre for Genomic Regulation (CRG), The Barcelona Institute of Science and Technology, Dr. Aiguader 88, Barcelona 08003, Spain; Wellcome Sanger Institute, Wellcome Genome Campus, Hinxton, Cambridge, UK; Universitat Pompeu Fabra (UPF), Barcelona, Spain; ICREA, Pg. Lluis Companys 23, Barcelona 08010, Spain

## Abstract

Many functions of proteins are performed by independently folding structural units called domains. The structures of domains are conserved during evolution but they are not identical. For example, the >270 human PDZ domains vary in the number of secondary structure elements and in the length of loops. An important but largely unexplored question is the impact of these extensions on protein energy landscapes: beyond any immediate functional effects, do extensions also alter the consequences of perturbations elsewhere in the domain, altering the potential for regulation and evolvability? Here we perform massively parallel energetic measurements on a model human PDZ domain to directly and comprehensively answer this question. In total we quantify the binding to a ligand and abundance of ∼190,000 protein variants to quantify free energy changes for mutations throughout the canonical domain fold and ∼7,000 energetic couplings between these mutations and the two domain extensions, both alone and in combination. We find that both extensions—one structured and one more dynamic—substantially and specifically re-shape the energy landscape of the domain, with the removal of an ɑ-helix altering the energetic consequences of 424 mutations in 54 sites on fold stability and 420 mutations in 56 sites on binding to a ligand. These changes to the energy landscape alter the effects of 330 allosteric mutations, including at solvent-accessible surface sites. Extending or pruning the domain therefore reshapes its energetic and allosteric landscape, adding and removing opportunities for the allosteric control of protein function.

## Introduction

Independently folding domains are often considered the functional and evolutionary structural units of proteins^1–5^. For example, the human genome encodes >8,000 distinct domain families with a median of two different domains per protein and many proteins having substantially more^1,4^. Individual domains from each family have conserved secondary structure topologies and folds, but they can also differ, with frequent amino acid (aa) insertions or deletions in loops and by the addition of secondary structure elements, particularly at their N– and C-termini^6,7^. These ‘domain extensions’ can have important functional consequences, for example altering protein stability or the affinity of binding to ligands^6–15^.

The impact of a domain extension could be direct, for example adding new ligand contacts to a binding interface, or it could be indirect, influencing function at another site in the protein. Such indirect or allosteric effects are not well understood and are difficult to predict^(16–19)^. In addition, a domain extension could potentially alter the consequences of perturbations elsewhere in the protein. For example, an extension might alter the energy landscape of a protein such that a distant allosteric regulatory site is strengthened or weakened. It is this question that we seek to address in this manuscript: how do domain extensions alter the effects of perturbations throughout a domain.

Perturbations to proteins include binding to other proteins, nucleic acids or small molecules, covalent post-translational modifications, and mutations. Mutations are particularly powerful experimental perturbations as they can be introduced at every site throughout a protein. This approach – often called deep mutational scanning – uses pooled libraries of variants and sequencing to quantify changes in variant frequencies during selection experiments. Coupled to selections for a particular protein property or function, mutational scanning can quantify the effects of thousands of different perturbations to a protein in a single pooled experiment^20^.

To quantify the impact of domain extensions using mutations, we ask whether adding or removing a domain extension changes the effect of each mutation elsewhere in the domain. For biophysical properties, this question is whether adding or removing a domain extension alters the energetic effect of a mutation. Formally, the question is whether each mutation is energetically coupled to loss or gain of the extension^21^. Energetic couplings (also called genetic interactions or epistasis) between a mutation and an extension might affect one or multiple properties of a protein, for example its stability or affinity of binding to a ligand.

At one extreme, domain extensions might be ‘modular’ changes, having functional consequences but effects that are not coupled to (i.e. are energetically additive with) perturbations elsewhere in the protein. At another extreme, domain extensions might be strongly energetically coupled to other sites, altering the effects of perturbations throughout the domain. The extent of this coupling will have important evolutionary and functional consequences. For example, strong energetic coupling between domain extensions and the rest of a domain could result in the emergence of new allosteric sites i.e. extension of the domain could change the sites elsewhere in the domain where mutations or other modifications alter the activity of the protein. The extension of a domain by structured or unstructured sequences could therefore create or remove regulatory sites. Similarly, strong energetic coupling could result in domain extensions creating and destroying opportunities for therapeutic targeting by the gain or loss of allosteric activity at accessible sites (^22,23^). Similarly, if a domain extension is energetically coupled to other sites in the domain it will by definition alter the consequences of mutations at those sites, so altering the potential for evolution i.e. evolvability^24,25^.

Recently, we^16,17,26,27^ and others^28–30^ have developed methods to comprehensively quantify the energetic effects of mutations throughout protein domains using massively parallel assays and model fitting^31^. Using different experimental selections it is possible to quantify the effects of mutations on different protein properties, including fold stability (quantified as the Gibbs free energy of folding, ΔG_f_) and binding to interaction partners (quantified as the Gibbs free energy of binding, ΔG_b_). These multiplexed experimental approaches present an opportunity to comprehensively quantify the impact of domain extensions on protein energy landscapes (Fig. 1).

**Figure 1:**
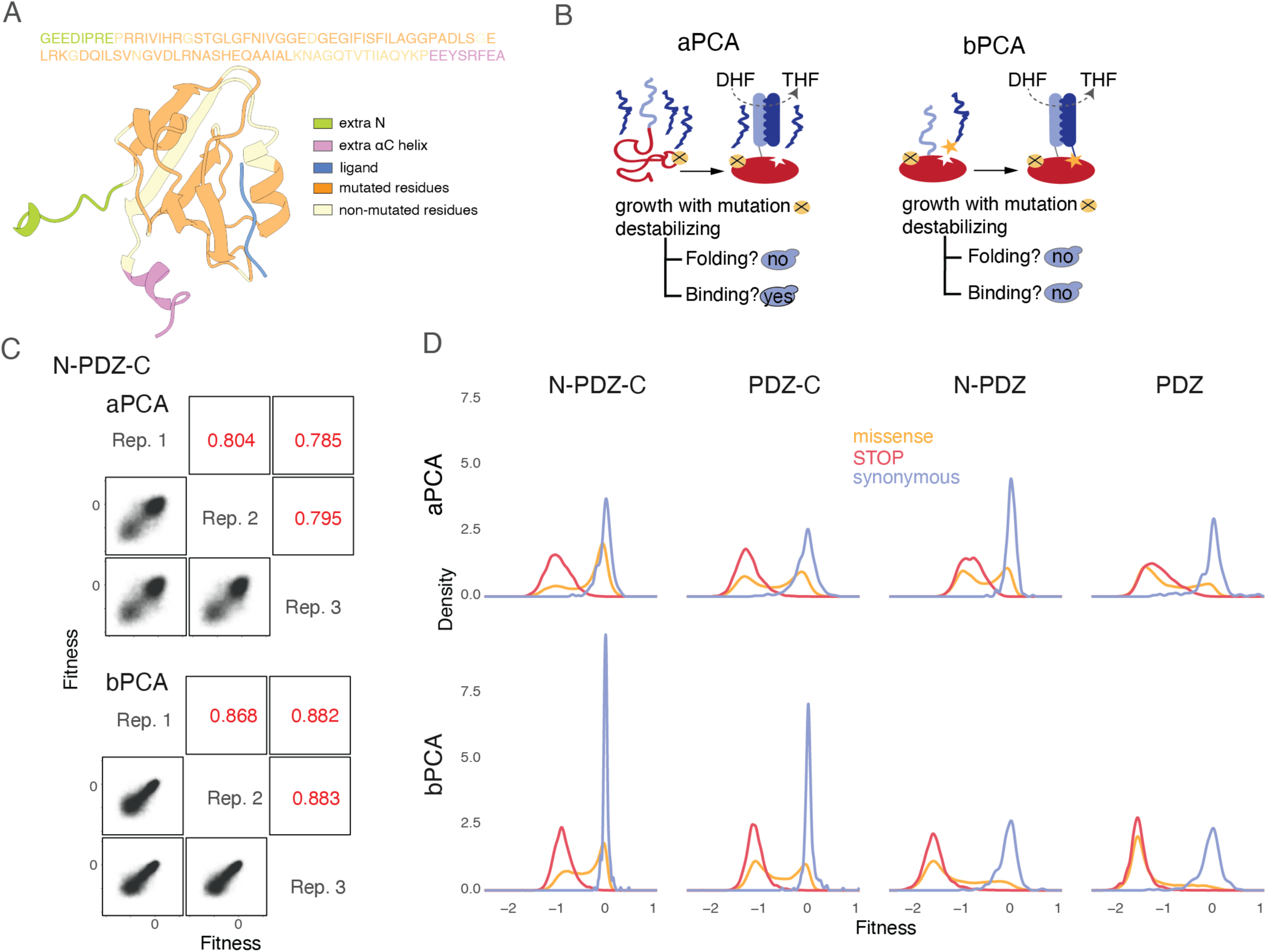
Quantifying the impact of protein domain extensions on the energy landscape of a protein domain. **a.** 3D structure and sequence of PDZ-CRIPT (adapted from PDB ID: 1BE9) indicating the N– and C-terminal extensions (green and violet respectively); the canonical fold (in orange the mutated and in pale yellow the non-mutated residues) and the CRIPT ligand (blue). **b.** Overview of AbundancePCA (aPCA) and BindingPCA (bPCA) selections. Yes, yeast growth: no, yeast growth defect; DHF, dihydrofolate; THF, tetrahydrofolate. **c.** Scatter plots showing the reproducibility of fitness estimates from aPCA and bPCA for the full-length domain (N-PDZ-C). Pearson’s R indicated in red. Rep., replicate. **d.** Fitness density distributions. Total variant counts for missense (yellow), premature STOPs (red) and synonymous mutations (blue) are indicated.

Here we apply this approach at scale. First, we measure the energetic effect of each mutation throughout a domain on fold stability (ΔΔG_f1_) and binding to a ligand (ΔΔG_b1_). We then repeat the measurements after removing the domain extension. The difference in the energetic effect of each mutation with and without the extension quantifies the energetic coupling between the extension and the mutation. For fold stability this coupling is quantified as ΔΔΔG_f1,2_ = ΔΔG_f1_ – ΔΔG_f2_, where ΔΔG_f2_ is the folding energy change of the mutation in the domain without the extension. For ligand binding it is quantified as ΔΔΔG_b1,2_ = ΔΔG_b1_ – ΔΔG_b2_, where ΔΔG_b2_ is the binding energy change without the extension (Fig. 2). These pairwise (second order) energetic couplings (genetic interactions) provide a complete picture of how removing a domain extension alters the energetic effects of mutations throughout a domain.

**Figure 2:**
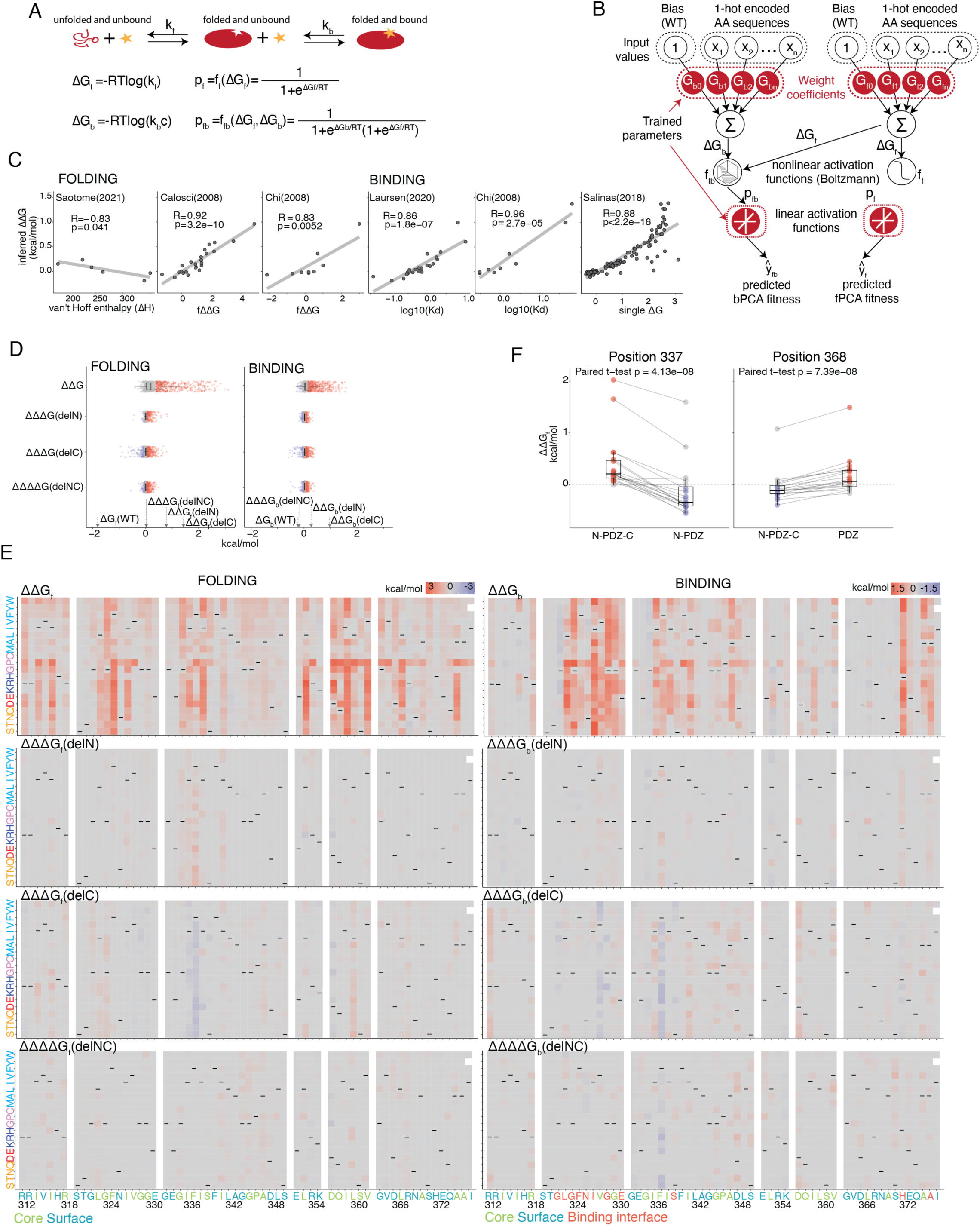
From molecular phenotypes to free energies. **a.** Three-state equilibrium and corresponding thermodynamic model. ΔG_f_, Gibbs free energy of folding; ΔG_b_, Gibbs free energy of binding; K_f_, folding equilibrium constant; K_b_, binding equilibrium constant; c, binding partner concentration; p_f_, fraction folded; p_fb_, fraction folded and bound; f_f_, nonlinear function of ΔG_f_; f_fb_, nonlinear function of ΔG_f_ and ΔG_b_; R, gas constant; T, temperature in Kelvin. **b.** Neural network architecture used to fit thermodynamic models to the PCA data (bottom, target and output data), thereby inferring the causal changes in free energy of folding and binding associated with single amino acid substitutions (top, input values). **c.** Comparisons of the model-inferred free energy changes to previously reported *in vitro* measurements^33,54–57^. Pearson’s R is shown. **d.** Energy values (kcal/mol) for first (ΔΔG), second (ΔΔΔG(delN), ΔΔΔG(delC)) and third (ΔΔΔΔG(delNC)) order energy terms for mutations for Folding and Binding phenotypes. Points are coloured depending on significance and sign of the mutation (grey: non-significant, red: significant and >0, blue: significant and <0) (Z-test, FDR < 0.05 and |inferred energy term|>median+MAD, see Materials and Methods). Boxplots show median and interquartile range only (no outliers). Arrows point to the kcal/mol values for global energy terms for WT, N-extension and C-extension deletions. **e.** Heatmaps showing inferred changes in folding and binding free energies (ΔΔG_f_ and ΔΔG_b_, top panels), changes in ΔΔG between full-length and truncated domains (ΔΔΔG_f_ and ΔΔΔG_b_, middle panels) and additional energetic contributions when both extensions are deleted together (ΔΔΔΔG_f_ and ΔΔΔΔG_b_, bottom panels). Amino acid labels in the x axis are coloured according to the solvent exposure in the full-length domain (green: rSASA<0.2, blue: rSASA>0.2. For Binding also residues in the binding interface (defined as residues with minimum side chain heavy atom distance to the ligand less than 5 Å) are coloured in red. **f.** Boxplots with paired data points (paired t-test performed) for Folding. Each dot represents the ΔΔG_f_ for each mutation of residue 337 (left) or 368 (right). Red (ΔΔG_f_>0) and blue (ΔΔG_f_<0) points are |ΔΔG_f_>0| (Z-test, FDR < 0.05). Lines connect the same mutations across datasets. Boxplots show median and interquartile range only (no outliers).

As a model system we use a well-studied model domain, the third PDZ domain from the protein PSD-95^16,25,32–35^ (henceforth referred to as PDZ3; PDZ domains are named for the three founding members of the family: Postsynaptic density protein (PSD-95), Drosophila discs large tumor suppressor (Dlg), and Zonula occludens-1 protein (ZO-1)^36^. PSD-95 is a member of the MAGUK (membrane-associated guanylate kinase) family and a key scaffolding protein found in excitatory synapses where it anchors proteins to the postsynaptic membrane^37^. PDZ domains are the largest family of human protein-protein interaction domains with more than 270 PDZ domains in >150 different human proteins that participate in a wide range of cellular processes, including signaling, cell polarity, cell adhesion, and neuronal synaptic transmission^38^. Many PDZ domains are considered important therapeutic targets, but very few have been successfully targeted, even experimentally^39,40^. PDZ domains bind diverse, normally C-terminal short peptide ligands. Despite low sequence identity, PDZ domains share a canonical fold composed of five to six β-strands and two or three α-helices. Ligand recognition occurs in a pocket composed of the β2 strand, the α2 helix, and the carboxylate-binding β1-β2 loop^41,42^ (Fig. 1a). Although PDZ domains share a conserved structural fold, they vary in the precise number of secondary structure elements and in the length of loops. These extensions are frequently at the N– and C-termini of the domains^6^.

PDZ3 from PSD-95 contains two domain extensions in addition to the canonical PDZ domain fold: an extra α-helix of nine aa (residues 394-402) at the C-terminus allied the α3 helix and a more dynamic N-terminal extension of eight aa (residues 303-310) (Fig. 1a)^6,43–45^. The α3 helix increases the affinity of PDZ3 binding to peptide ligands ∼20-fold^44^. The α3 helix does not directly contact the ligand, so the change in binding affinity must have an allosteric mechanism. The allosteric potential of the α3 helix is further demonstrated by the consequences of phosphorylation of Y397 within the helix: this post-translational modification further increases ligand binding affinity^446^. Engineering a photoconvertible residue into the α3 helix further and elegantly demonstrated its allosteric impact on binding affinity^47^. The impact of the N-terminal extension to PDZ3 is less studied^48^.

Here we use an experimental design that allows us to comprehensively quantify the changes in folding energy and ligand binding energy for mutations throughout the PDZ3 domain in four different contexts: in the presence of both domain extensions, in the absence of the α3 helix, in the absence of the N-terminal extension, and in the absence of both extensions (Fig. 1a). In total we quantify 187,622 mutational effects on protein abundance and 195,047 mutational effects on ligand binding to quantify the energetic couplings between all aa substitutions and deletion of each of the N– and C-terminal extensions, as well as the energetic interactions with deletion of both extensions together. Both extensions alter the stability of the domain and its ligand binding affinity. In addition, both extensions are also energetically coupled to a subset of sites throughout the core PDZ domain, quantifying how the extensions modulate the core domain’s folding and binding energy landscapes. One consequence of this is a transformation of the domain’s allosteric landscape, altering evolvability and the potential for regulation and therapeutic targeting.

## Results

### Domain extensions in PSD-95 PDZ3

PDZ domains share a common fold consisting of five to six β-strands with two to three α-helices and bind peptide ligands in a groove formed between the second β-strand and the second α-helix^41,42^. PSD-95 (postsynaptic density protein 95, encoded by the *DLG4* gene) is a major scaffolding protein at the postsynaptic density of excitatory synapses and contains three PDZ domains. The third PDZ domain of PSD-95, henceforth PDZ3, is one of the best-characterized PDZ domains^25,32–35^ but contains two domain extensions: an additional short six aa C-terminal helix, αC, that packs against the β-sandwich core of the PDZ domain, and a more dynamic eight aa N-terminal extension (henceforth, N-extension) that contacts the first two β-strands on the back side of the domain relative to the ligand-binding site^6^ (Fig. 1a).

To comprehensively characterise the impact of these extensions on the energetic landscape of PDZ3, we designed an experiment to quantify the changes in protein stability (Gibbs free energy of folding, ΔG_f_) and ligand binding (Gibbs free energy of binding, ΔG_b_) for mutations throughout the canonical fold of the PDZ domain in four different contexts: [1] the full length PDZ3 domain containing both extensions (N-PDZ-C, aa 303 to 402), [2] PDZ3 without the N-extension (PDZ-C, aa 311 to 402), [3] PDZ3 without the αC helix (N-PDZ, aa 303 to 394), and [4] PDZ3 without both extensions (PDZ, aa 311 to 394) (Fig. 1a).

In each context we constructed a library of single and double aa variants covering 66 aa of the canonical PDZ domain (residues 312 to 377). Each library was constructed by co-mutating 144 pairs of sites with all 19 aa variants in each position, creating a designed diversity of 53,144 genotypes (51,984 double mutants and 1,159 single mutants, see Methods). In each of the four contexts, the effect of each aa substitution is therefore tested individually and in a large number of double mutants. Quantifying the effect of each mutation in a large number of double mutants allows inference of the underlying causal energetic effects of mutations and their energetic interactions with the domain extensions by thermodynamic model fitting^16,31,49^.

### Quantifying abundance and ligand binding for ∼190,000 genotypes across four extension contexts

For each genotype in each of the four contexts we quantified two molecular phenotypes using pooled selection experiments and deep sequencing. In the first experiment, we used a highly validated abundance protein-fragment complementation assay (AbundancePCA, aPCA;^16,27,50,51^) to quantify the concentration of folded protein for each variant inside yeast cells. In the second assay, we used BindingPCA (bPCA;^16,35,4116,50,52^^16,35,41,53^) to quantify binding to a ligand: a canonical class I C-terminal peptide ligand from the protein CRIPT^16,25,32,35,41^.

All experiments were performed in three biological replicates, necessitating a total of 24 pooled selection-sequencing experiments (four contexts x two phenotypes x three replicates).

Abundance and binding measurements were highly reproducible (median Pearson correlation coefficient, *r* = 0.815 and *r* = 0.844 for aPCA and bPCA, respectively; Fig. 1c and Extended Data Fig. 1b). The results also correlated well with previous measurements for the canonical domain without domain extensions (Pearson correlation coefficient *r* = 0.86, n=1548 and *r* = 0.89, n=1883 for aPCA and bPCA, respectively, Extended Data Fig. 1c).

Abundance and binding fitness values for all experiments were bimodally distributed, with the upper and lower modes centred on the fitness values of synonymous and stop variants, respectively (Fig. 1d). However, the proportion of variants in the lower fitness peak differed across the experiments, with the smallest proportion of detrimental genotypes in the N-PDZ-C library, followed by PDZ-C, N-PDZ, and the largest number of detrimental variants in the shortest PDZ construct lacking both extensions ( Fig. 1d). This is consistent with both the N– and C-extensions stabilising the domain, with a larger effect of the C-extension.

### From molecular phenotypes to free energy changes

We used the comprehensive measurements of abundance and binding in double mutants to infer the underlying causal changes in the free energy of folding (ΔG_f_) and binding (ΔG_b_). We used MoCHI^31^ to fit a single thermodynamic model to the entire dataset (Extended Data Fig. 2c and Materials and Methods). In this model, the fraction of the protein that is folded (and so aPCA fitness) depends non-linearly on the Gibbs free energy of folding (ΔG_f_). In contrast, the fraction of the protein bound to the ligand (and so bPCA fitness) depends non-linearly on both the folding energy (ΔG_f_) and the binding energy (ΔG_b_), calculated using the Boltzmann distribution partition function (Fig. 2a). The model assumes that each mutation and domain extension has a fixed effect on the folding (ΔΔG_f_) and binding (ΔΔG_b_) energy, quantified in the full-length domain (N-PDZ-C). However the model also allows pairwise energetic couplings (ΔΔΔG_f_ and ΔΔΔG_b_) between all mutations, including between each substitution and the deletion of each of the two domain extensions. In addition, the model also allows for third order interactions between each aa substitution, the N-extension deletion, and the C-extension deletion (ΔΔΔΔG_f_ and ΔΔΔΔG_b_, see methods). These third order energy terms quantify any additional synergistic or antagonistic effects when both extensions are deleted together.

The abundance and binding fitness of each genotype is calculated by summing the free energy changes and transforming the total energy to molecular phenotypes using the Boltzman partition functions. For example, the binding free energy of a double mutant containing two mutations A and B and a C-terminal extension deletion (ΔG_b_(N-PDZ,A,B)) is modelled as the sum of the wild-type free energy (ΔG_b_(N-PDZ-C)), the changes in free energy for each of the 3 mutations (ΔΔG_b_ for the A and B mutations, and ΔΔG_b_ for the deletion of the ɑC helix, referred to as the delC mutation) and three ΔΔΔG_b_ pairwise energetic couplings (ΔΔΔG_b_(A,delC), ΔΔΔG_b_(B,delC) and ΔΔΔG_b_(A,B)):

ΔG_b_(N-PDZ,A,B) = ΔG_b_(N-PDZ-C) + ΔΔG_b_(A) + ΔΔG_b_(B) + ΔΔG_b_(delC) + ΔΔΔG_b_(A,delC) + ΔΔΔG_b_(B,delC) + ΔΔΔG_b_(A,B)

The model provides excellent predictive performance (mean of the variance explained R^2^ = 0.73 and R^2^ = 0.75, evaluated by 10-fold cross-validation for abundance and binding, respectively, Extended Data Fig. 2d) and the inferred free energy changes agree very well with previous in vivo^16^ (*r* = 0.74 *r* = 0.59 for Folding and Binding respectively, Extended Data fig. 2e) and in vitro measurements^33,54–57^ (mean Pearson correlation coefficient *r* = 0.88 (range 0.83 to 0.96), Fig. 2c).

### Overview of the folding and binding energy landscapes

We first considered the individual energetic effects of the domain extensions. Consistent with previous experiments^43,44^, removing the αC helix destabilizes the domain (ΔΔG_f_(delC) = 1.41 kcal/mol) and allosterically reduces the ligand binding affinity (ΔΔG_b_(delC) = 0.97 kcal/mol) (arrows in Fig. 2d). Removing the N-extension also reduces the fold stability, although to a lesser extent (ΔΔG_f_(delN) = 0.78 kcal/mol) and causes a small reduction in binding affinity (ΔΔG_b_(delN) = 0.27 kcal/mol). Combined deletion of both the αC helix and the N-extension has an additive effect on stability (interaction energy, ΔΔΔG_f_(delNC) = 0.011) but the reduction in binding affinity is less than the sum of the individual deletions (ΔΔΔG_b_(delNC) = –0.21 kcal/mol), an example of positively epistasis.

We next considered the individual effects of the substitutions in the PDZ domain (ΔΔG_f_ and ΔΔG_b_ in the full-length N-PDZ-C construct), how they interact with removal of each extension (ΔΔΔG_f_(delN), ΔΔΔG_f_(delC), ΔΔΔG_b_(delN), ΔΔΔG_b_(delC)), and how they interact with the combined deletion of both extensions (ΔΔΔΔG_f_(delNC), ΔΔΔΔG_b_(delNC)). In the heatmaps in Fig. 2e we plot all eight energy terms for all 1158 mutations. The first order individual effects of the mutations on folding (ΔΔG_f_) and binding energy (ΔΔG_b_) are typically much larger than the six interaction energies (ΔΔΔG_f_(delN), ΔΔΔG_f_(delC), ΔΔΔΔG_f_(delNC), ΔΔΔG_b_(delN), ΔΔΔG_b_(delC), ΔΔΔΔG_b_(delNC)) (Fig. 2d). The first order energies have distributions typical for a small globular protein, with large changes in ΔΔG_f_ for mutations in residues buried in the hydrophobic core (t-test, p ≤ 0.0001, Extended Data Fig. 2a) and large changes in ΔΔG_b_ for mutations in many (but not all) residues that directly contact the ligand (t-test, p ≤ 0.0001,Extended Data Fig. 2a). In the full-length domain, 411 mutations in 53 sites change the fold stability (FDR<0.05, Z-test and |ΔΔG_f_|>(median ΔΔG_b_+Median Absolute Deviation(MAD)) and 438 mutations in 48 sites change the binding affinity (FDR<0.05 and |ΔΔG_b_|>(median ΔΔG_b_ +MAD)).

The interaction energies are, however, different, with most mutations in most sites having interaction energies very close to zero (Fig. 2d). A subset of sites are, however, energetically coupled to the domain extensions (Fig. 3b and Fig. 5b)

**Figure 3:**
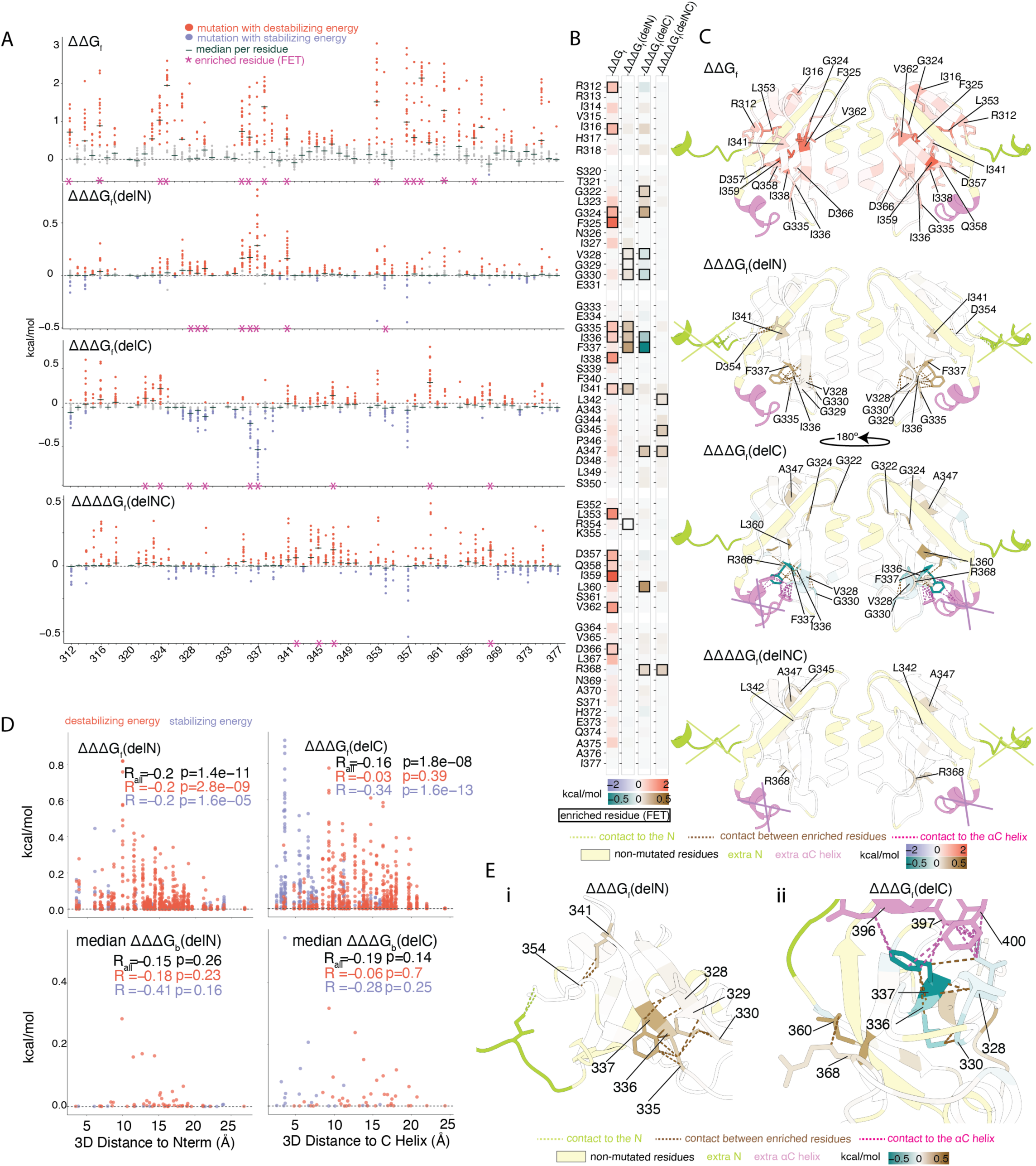
Impact of domain extensions on folding energies. **a.** Manhattan plots showing the folding energy changes of all single aa substitutions (ΔΔG_f_, top panel, ΔΔΔG_f_, middle panels and ΔΔΔΔG_f_, bottom panel). Points are coloured depending on significance and sign of the mutation (grey: non-significant, red: significant and >0, blue: significant and <0) (Z-test, FDR < 0.05 and |inferred energy term|>median+MAD, see Materials and Methods).The dash indicates the per-residue median energy values. Asterisks indicate the residues that are enriched in significant mutations (FET, FDR<0.1). **b.** Heatmaps showing the median energy value per residue. Bold squares indicate residues enriched for significant mutations (FET, FDR<0.1). **c.** Structures of the PDZ domain (adapted from PDB ID: 1BE9) coloured by the per-residue median (top: ΔΔG_f_, middle: ΔΔΔG_f_(delN) and ΔΔΔG_f_(delC), bottom: ΔΔΔΔG_f_(delNC)) (non-mutated residues in pale yellow). Residues enriched in significant mutations are indicated in black. Deleted extensions (N in green and C in violet) are indicated. Brown dashed lines connect contacting residues. Green and violet dash lines represent contacts between residues and N or C respectively. **d.** Relationship between |ΔΔΔG_f_(delC)| and the distance to the αC helix (right) and |ΔΔΔG_f_(delN)| and the distance to the N (left) (minimal side chain heavy atom distance). Upper panels show the energies for each mutation and the bottom panel show the per-residue median. Points are coloured as in a. Pearson’s R for all the points is in black, for positive energies are in red and for negative energies are in blue. **e.** Zoom view of contacting residues from c.

**Figure 4:**
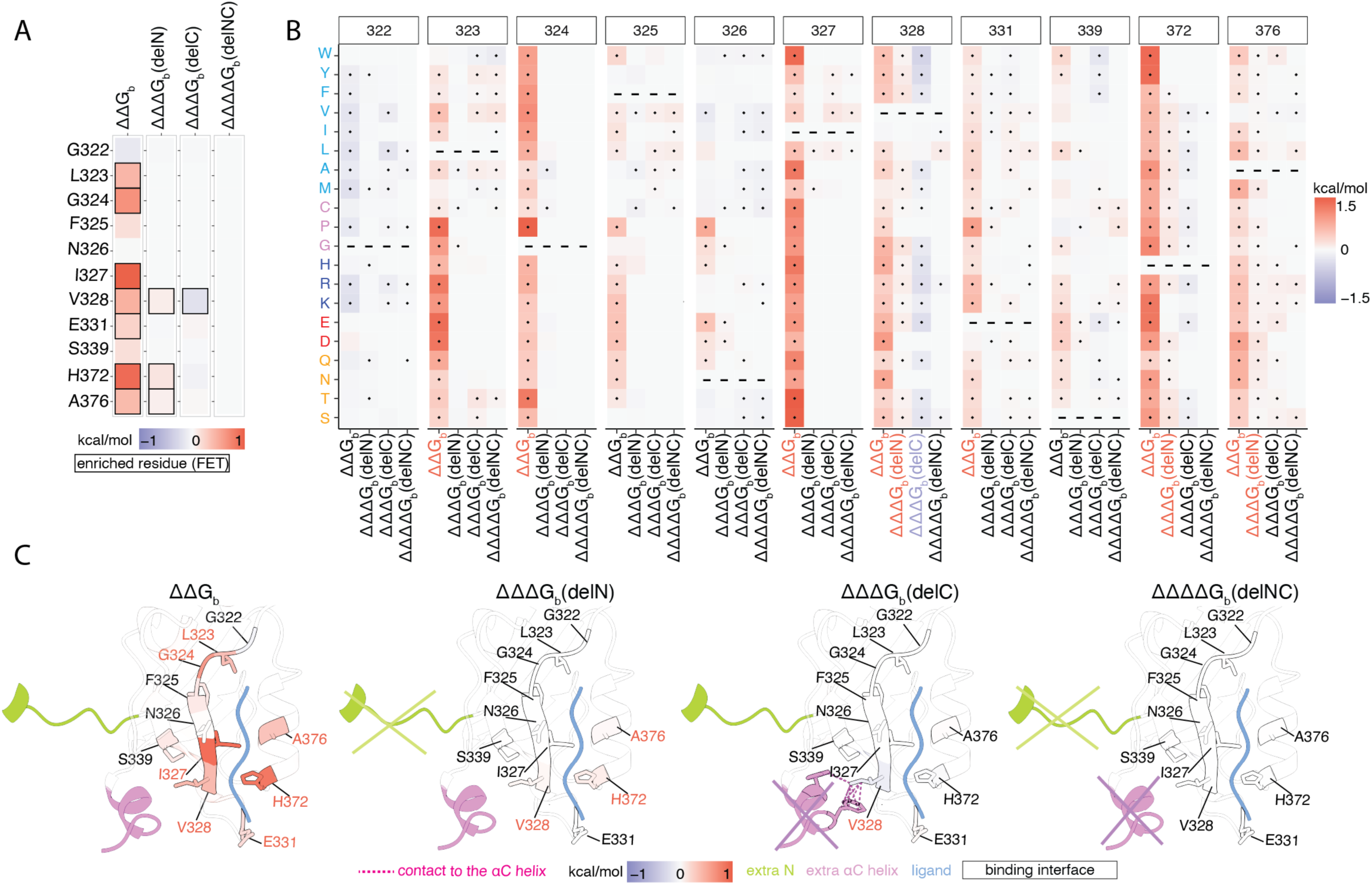
Impact of domain extensions on binding energies in the binding interface. **a.** Heatmaps of the residues in the binding interface (minimal side chain heavy atom distance to ligand) showing the median energy value per residue. Bold squares indicate residues enriched for significant mutations (FET, FDR<0.1). **b.** Heatmaps showing inferred changes in binding free energies (ΔΔG_b_, first column for each residue), changes in ΔΔG between full-length and truncated domains (ΔΔΔG_b_(delN) and ΔΔΔG_b_(delC), second and third columns for each residue respectively) and additional energetic contributions when both extensions are deleted together (ΔΔΔΔG_b_(delNC), last column for each residue). Colours in x axis labels indicate enrichment for significant mutations (FET, FDR<0.1) and sign of energy change (black for non-enriched residues, red for enriched residues with positive energy change and blue for enriched residues and negative energy change). Dots are significant mutations (Z-test, FDR < 0.05 and |inferred energy term|>median+MAD) **c.** Structures of the PDZ domain (adapted from PDB ID: 1BE9) coloured by the median energy value per residue (left: ΔΔG_f_, middle: ΔΔΔG_f_(delN) and ΔΔΔG_f_(delC), right: ΔΔΔΔG_f_(delNC)) (non-mutated residues in pale yellow). Binding interface residues are labeled and highlighted in black. Residues enriched in significant mutations are indicated in red. Deleted extensions (N in green and αC in violet) are indicated.

**Figure 5:**
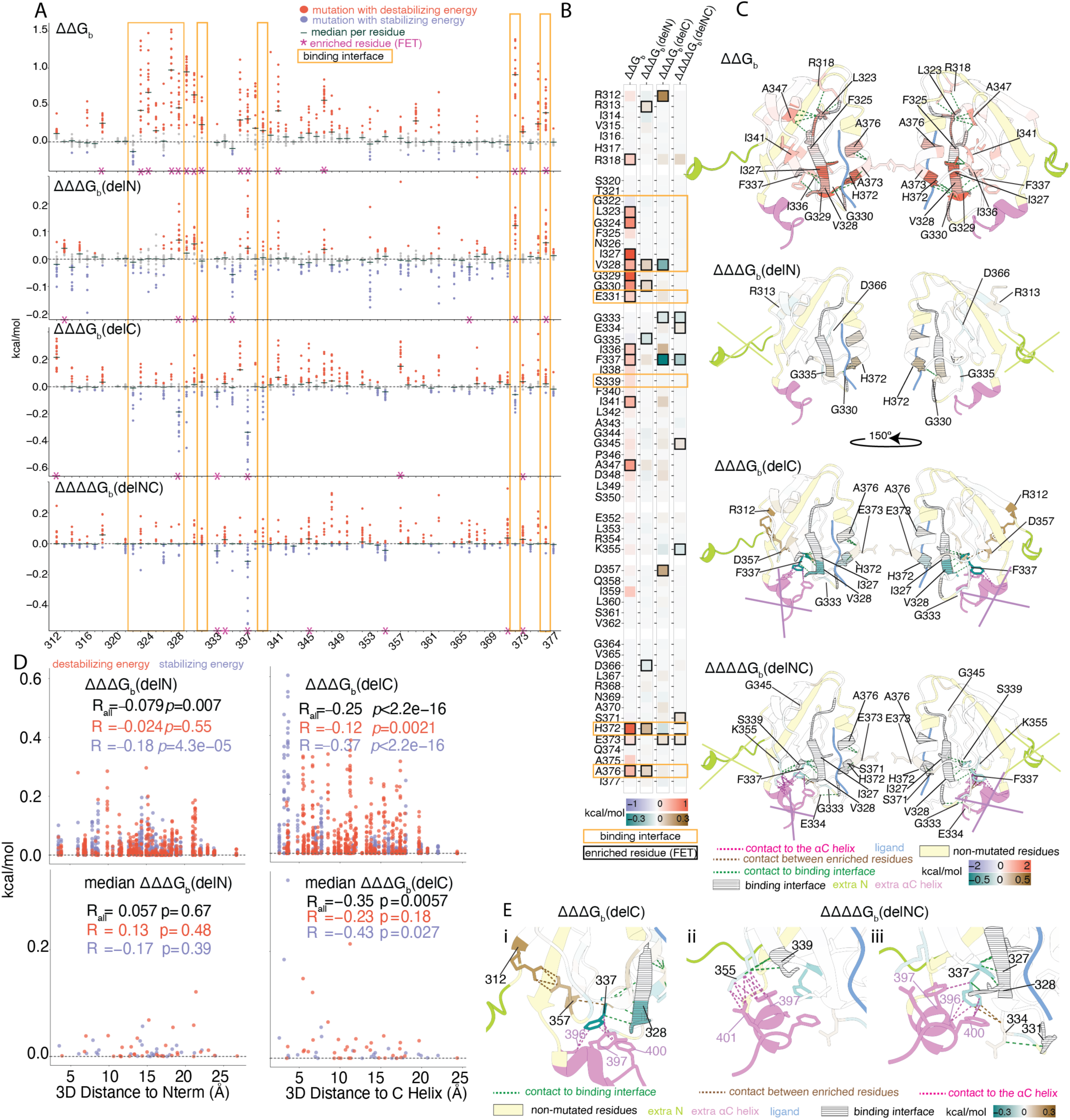
Changes in binding energies outside the binding interface. **a.** Manhattan plots showing the binding energy changes of all single aa substitutions (ΔΔG_b_, top panel, ΔΔΔG_b_, middle panels and ΔΔΔΔG_b_, bottom panel). Points are coloured depending on significance and sign of the mutation (grey: non-significant, red: significant and >0, blue: significant and <0) (Z-test, FDR < 0.05 and |inferred energy term|>median+MAD, see Materials and Methods).The dash indicates the per-residue median energy values. Asterisks indicate the residues that are enriched in significant mutations (FET, FDR<0.1). **b.** Heatmaps showing the per-residue median energy value. Bold squares indicate residues enriched for significant mutations (FET, FDR<0.1). **c.** Structures of the PDZ domain (adapted from PDB ID: 1BE9) coloured by the per-residue median (top: ΔΔG_b_, middle: ΔΔΔG_b_(delN) and ΔΔΔG_b_(delC), bottom: ΔΔΔΔG_b_(delNC)) (non-mutated residues in pale yellow). Residues enriched in significant mutations are indicated in black. Deleted extensions (N in green and C in violet) are indicated. Brown dashed lines connect contacting residues. Dark green, light green and violet dash lines represent contacts between residues and the binding interface, N or αC respectively. The binding interface is filled with black dashed lines. **e.** Relationship between |ΔΔΔG_b_(delC)| and the distance to the αC helix (right) and |ΔΔΔG_b_(delN)| and the distance to the N (left) (minimal side chain heavy atom distance). Upper panels show the energies for each mutation and the bottom panel show the median of the energy values per residue. Points are coloured as in a. Pearson’s R for all the points is in black, for positive energies are in red and for negative energies are in blue. **d.** Zoom view of contacting residues from c.

In total, for fold stability, eight positions are enriched for mutations energetically coupled to the N-extension, nine are enriched for mutations energetically coupled to the αC extension, and four are enriched for mutations energetically coupled to the combination of both extensions (FDR<0.1 Fisher’s exact test (FET)); residues marked with an asterisk in Fig. 3a and bold squares in Fig. 3b).

For binding, seven positions are enriched for mutations energetically coupled to the N-extension, six are enriched for mutations coupled to the αC extension, and seven are enriched for mutations coupled to the combination of both extensions (FDR<0.1 FET, residues marked with an asterisk in Fig. 5a and squares in Fig. 5b). The mutational effects with and without the domain extensions differ mostly in magnitude. However, for mutations in two sites they also differ in sign, with mutations switching between being destabilizing and stabilizing in the different proteins, an example of sign epistasis^58,59^ (Extended Data Fig. 2b).

In the first of these examples, mutations at position 337 are destabilizing in the full-length protein (median ΔΔG_f_(N-PDZ-C) = 0.21 kcal/mol) but they have a negative interaction energy with deletion of the αC helix (median ΔΔΔG_f_(delC) = –0.55 kcal/mol). The effect of these mutations on folding in the shorter N-PDZ domain without αC is thus reversed, with the mutations now stabilizing the domain (median ΔΔG_f_(N-PDZ) = ΔΔG_f_(N-PDZ-C) + ΔΔΔG_f_(delC) = 0.21 + (–0.55) = –0.34 kcal/mol). Sign epistasis in the opposite direction happens for mutations at position 368, which have a median ΔΔG_f_(N-PDZ-C) = –0.12 kcal/mol) and a positive interaction energy with deletion of the αC helix (median ΔΔΔG_f_(delC) = 0.09 kcal/mol) and a positive third order interaction energy with deletion of both N and αC helix (median ΔΔΔΔG_f_(delNC) = 0.12 kcal/mol). The effect of these mutations on folding in the shorter PDZ domain without any extension is thus reversed, with mutations now reducing fold stability (median ΔΔG_f_(PDZ) = ΔΔG_f_(N-PDZ-C) + ΔΔΔG_f_(delC)+ΔΔΔG_f_(delN)+ΔΔΔΔG_f_(delNC) = (–0.12) + 0.09 + (–0.001) + 0.12 = 0.09 kcal/mol) (Extended Data Fig. 2b). Comparing ΔΔG_f_ for individual mutations for each residue shows that this change is significant (Fig. 2f, paired t-test).

### Domain extensions modulate the fold stability landscape

We examined the structural location of sites energetically coupled to each domain extension. When considering folding energies, the eight sites enriched for mutations coupled to the N-extension are located in the third β-strand (336, 337, 341), in the second loop (330, 335) that precedes this strand, in the second β-strand before it (328,329) and in the fourth loop (354). One of these residues (354) directly contacts the N-extension (position 307 in the extension). Position 354 also contacts position 341, which is in the same strand as 337 and 336, and the last two directly contact position 335 and each other with residue 336 also contacting 330, residue 337 also contacting 328 and residue 335 also contacting 329 (Fig. 3c,e). The residues with strongest energetically coupling to the N-extension thus form a spatially clustered network of structural contacts connected to the extension (Fig. 3(i)). However, they are not all close to the extension (Fig. 3d).

In contrast, the nine sites enriched for mutations energetically coupled to the αC helix are located in three spatial clusters. The first cluster consists of four residues where mutations typically have milder effects when the αC helix is deleted (ΔΔΔG_f_(delC)<0): 328 in the second strand and 330 in the second loop just after it, and 336 and 337 in the third strand. Phenylalanine 337 and valine 328 both directly contact each other and the αC helix as part of a hydrophobic interface with αC helix residues 396, 397 and 400 (Fig. 3e(ii)). Position 336 does not directly contact the αC helix but does contact position 337. In turn, position 330 contacts 336 (Fig. 3c,e). The second cluster of three residues contains two in the carboxylate-binding loop (322 and 324) and one in the second helix (347). Mutations in these three residues typically have stronger effects in the absence of the αC helix (ΔΔΔG_f_(delC)>0) as do mutations in the two residues of the third cluster: 360 in the fourth strand and 368 in the fifth loop (Fig. 3c). The side chains of residues 360 and 368 are solvent-exposed and contacting directly. Plotting the energetic couplings to the αC helix against the shortest Euclidean distance to it reveals that negative energetic couplings (mutations that have weaker effects on folding when the helix is removed) but not positive couplings tend to be located closer to the αC helix (Fig. 3d).

Positions enriched for third order energetic couplings with the removal of both the N-extension and the αC helix are a cluster of three residues – leucine 342 and glycine 345 in the third loop, alanine 347 in the first helix – and an additional residue, arginine 368 in the fifth loop (Fig. 3c). None of these residues directly contact either of the two extensions.

In summary, removing either or both domain extensions alters the effects of mutations on fold stability in a small subset of spatially clustered residues. 13% (8 out of 61) and 15% (9 out of 61) of sites are enriched for these energetically-coupled mutations for the N– and C-extensions, with 7% of sites (4 out of 61) enriched for mutations with effects that further alter when both extensions are removed. However, only two of these sites directly contact the αC helix (328 and 337) and only one directly contacts the N-extension (354). The rest of the changes in mutational effects must, therefore, have indirect—i.e. allosteric— mechanisms.

### Impact of domain extensions on the binding interface

We next asked how the domain extensions alter the energetic landscape of the PDZ3 ligand-binding interface. PSD-95 PDZ3 binds the CRIPT peptide in a groove between the second β-strand and the second α-helix, with the C-terminal carboxylate of the ligand recognized through backbone interactions within the β1-β2 loop^25,41^. In total, 11 PDZ residues are part of the binding interface (distance from sidechain heavy atom to the nearest ligand residue < 5 Å). Consistent with the concept of binding interface hotspots^60,61^), mutations in only a subset of the sites that contact the ligand cause a large change in the binding energy: only 7/11 residues that contact the ligand are enriched for mutations that change the binding energy in the full-length domain (FDR<0.1, FET, Fig. 4a).

Three interface positions are enriched for mutations with binding energy energetic couplings to the N-extension (328, 372 and 376), and one is enriched for mutations with binding energy energetic couplings to the αC helix (328). None is enriched for energetic couplings with the removal of both extensions (Fig. 4a). Positions 328, 372 and 376 do not directly contact the N-extension but 328 does contact the αC helix at positions 397 and 400 (Fig. 4c).

Both domain extensions therefore specifically modulate the effects of mutations in the binding interface, including by indirect allosteric effects.

### Domain extensions modulate the allosteric landscape

We next considered changes in binding energy due to mutations outside of the binding interface. Changes in binding energy caused by mutations in residues that do not contact the ligand must have an indirect—i.e. allosteric—mechanism^16–18^. In total, eight residues are enriched for allosteric mutations in the full-length PDZ domain (FDR<0.1, FET, Fig. 5b). These residues, which we refer to as major allosteric sites^16^, are located throughout the PDZ domain. Residue 318 in the first loop contacts position 323 in the binding interface residue 329 at the end of the second strand and residue 330 in the second loop both contact position 372 in the binding interface; in the third strand, residue 336 contacts position 327 in the interface and residue 337 contacts position 328 in the interface; residue 347 in the first helix contacts positions 323 and 325 in the interface; and residue 373 in the second helix contacts residues 372 and 376 in the interface (Fig. 5c). In total, therefore, seven out of eight sites enriched for allosteric mutations are second shell residues that directly contact at least one residue in the binding interface; only residue 341 does not.

Removing each domain extension changes a subset of the allosteric mutational effects. In total, four sites outside of the binding interface are enriched for mutations energetically coupled to loss of the N-extension (FDR<0.1, FET), five are enriched for mutations energetically coupled to loss of the αC helix (FDR<0.1, FET), and seven are enriched for mutations whose effects on binding energy further change when both the N-extension and the αC helix are removed (FDR<0.1, FET, although note the third order energetic couplings are very small in magnitude, Fig. 5a,b).

Plotting the energetic couplings for each mutation and the median coupling for each site (Fig. 5d) against the distance from each extension reveals that mutations with weaker effects when the αC helix is removed (ΔΔΔG_b_(delC)<0) tend to be located closer to the αC helix, but this is not true for mutations with stronger effects or mutations coupled to removal of the N-extension (Fig. 5d).

None of the four sites enriched for mutations with binding energy couplings to the N-extension are in contact with it, although residue 313 located in the first β-strand is close in the linear sequence. Moreover, only position 330 is in direct contact with a residue in the binding interface (contacting position 372). Positions 330 and 335 are both glycine residues in the second loop of the domain. Positions 313 and 366 are far from the binding site, both with side chains facing the solvent (Fig. 5c).

Of the five sites enriched for mutations with binding energy couplings to the αC helix, position 337 directly contacts both the αC helix (residues 396, 397 and 400) and the binding interface (position 328, which is also enriched for mutations coupled to the αC helix and to position 327) (Fig. 5c). Position 337 directly contacts residue 357 that in turn contacts 312 through a salt bridge (Fig. 5c,e(i)). Residue 373 contacts position 372 and 376 in the binding interface (Fig. 5c). Three of the five sites where mutational effects on binding energy change when the αC helix is deleted outside the binding interface therefore form a continuous network connected to both the αC helix and the binding interface (Fig. 5c,e(i)), and another residue directly contacts the binding interface.

Similarly of the seven residues enriched for mutations with third-order binding energy couplings to removal of both extensions, position 355 in the fourth loop and 337 in the third β-strand directly contact both the αC helix (355 contacting residues 397 and 401 and 337 contacting residues 396,397 and 400) and the binding interface (355 contacting 339 and 337 contacting 328 and 327) (Fig. 5c,e(ii)). Positions 371 and 373 contact the binding interface residue 372 (373 contacts also 376 in binding interface). Position 334 contacts directly with 337 and also with 331 in the binding interface. (Fig. 5c,e(iii)). The other two residues are glycines located in the second (333) and third (345) loops and do not contact either extension nor the binding interface. However residue 333 is next to residue 334 in the linear sequence and the β-strand that both loops are flanking contains position 337, which is also enriched and in contact with both the αC helix and the binding interface, and position 339, which is in the binding interface. Three of the seven residues energetically coupled to removal of both extensions form a continuous network connecting the αC helix and the binding interface and two residues contact the binding interface directly.

### Changes in allosteric hotspots

In PDZ3—as in other allosteric proteins^16–18^—mutations in residues closer to the interface have larger effects on the binding energy (Fig. 6a and Extended Data Fig. 3). This distance-dependent allosteric decay is observed in all complete allosteric maps generated to date and appears to be a conserved principle of protein biophysics^16–18,62–64^.

**Figure 6:**
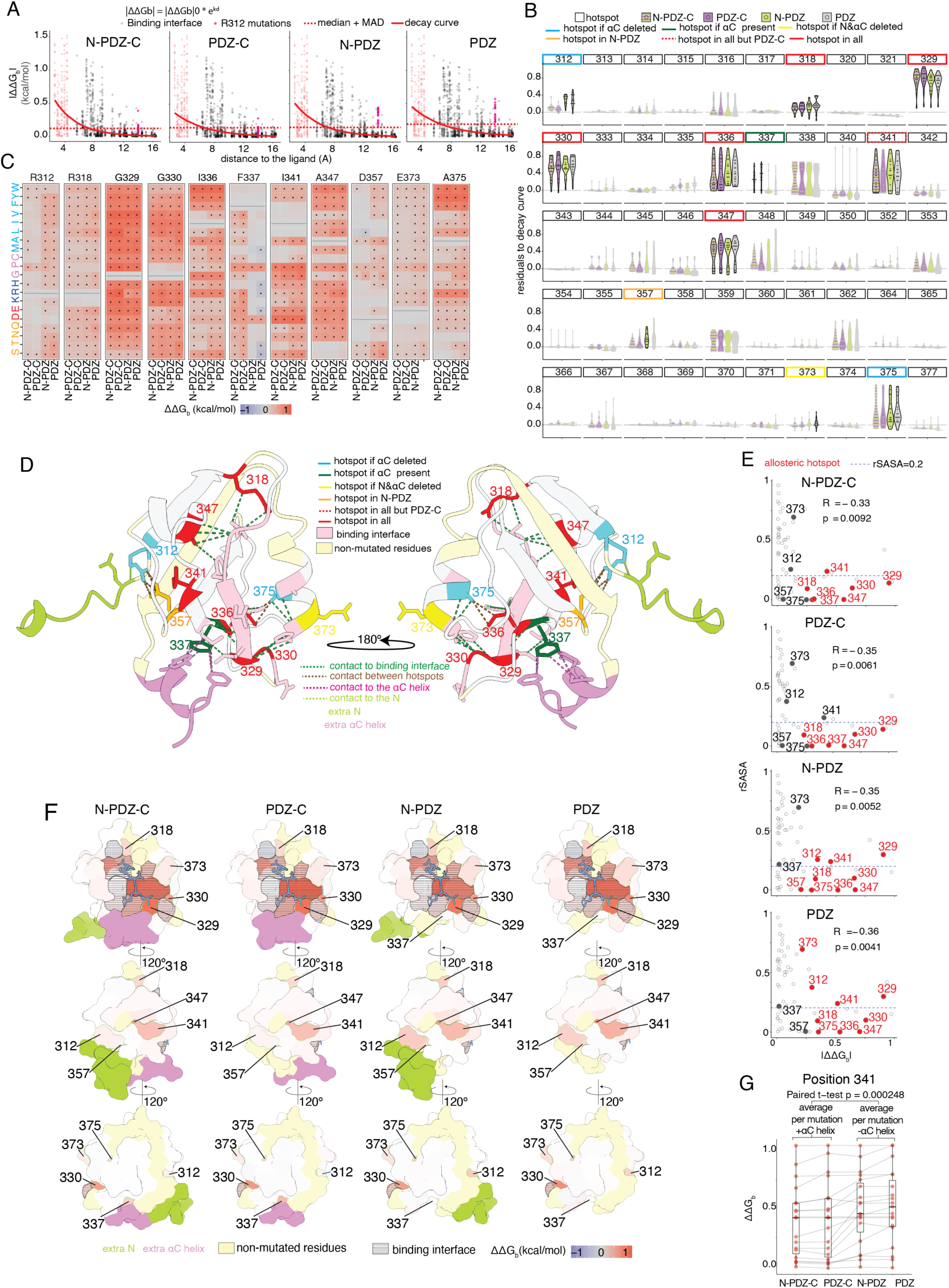
Transformation of the allosteric landscape. **a.** Exponential decay fits to the relationship between |ΔΔG_b_| to the minimum heavy atom distance to the ligand. |ΔΔG_b_|_0_=starting ΔΔG_b_ at distance d = 0, k = decay rate, d = distance from the ligand. Points are red if in the binding interface (minimal side chain heavy atom distance to ligand < 5 Å). Magenta points correspond to mutations of the residue 312. Dash red line is in |ΔΔG_b_| = median + MAD, see Materials and Methods. **b.** Violin plots showing the residuals to the decay curves in **a** for each of the 50 mutated residues outside the binding interface and for each of the four PDZ domain lengths (N-PDZ-C in violet and green stripes, PDZ-C in violet, N-PDZ in green and PDZ in grey). Residue numbers squares are coloured depending on being hotspots for all the four domains (red), three domains (dashed red), only if αC is deleted (cyan), only if αC is present (dark green), only in N-PDZ (orange) and only if N and αC are deleted (yellow). **c.** Heatmaps showing ΔΔG_b_ for each of the 11 allosteric hotspots and for each of the four domains. Dots are significant mutations (Z-test, FDR < 0.05 and |ΔΔG_b_|>median+MAD) **d.** Structures of the PDZ domain (adapted from PDB ID: 1BE9). Residues are coloured as in **b**. Brown dashed lines connect contacting hotspots. Dark green, light green and violet dash lines represent contacts between hotspots and binding interface, N or αC respectively. **e.** Relationship between the position-wise median change in free energy of binding |ΔΔG_b_| and the solvent exposure of the corresponding residue (rSASA). R = Pearson’s coefficient. The eleven hotspots are labelled (red if hotspot in that domain). The blue dashed line is at rSASA = 0.2. **f.** Surfaces (adapted from PDB ID: 1BE9) are coloured by the per-residue median ΔΔG_b_ (non-mutated residues in pale yellow). The binding interface is filled with black dashed lines. The eleven hotspots are labeled. The ligand is coloured in blue. **g.** Comparison of mutational ΔΔG_b_ at position 341 across the four domains. Each dot represents a mutation, and dots belonging to the same mutation are connected across datasets by thin lines. Boxplots indicate the distribution of ΔΔG_b_ within each domain, displaying median and interquartile range only. p-value from a paired t-test comparing the average per-mutation energy of N-PDZ-C and PDZ-C (containing αC helix group) vs N-PDZ and PDZ (without αC helix group) is shown.

However, mutations in a subset of sites have allosteric effects that are stronger than expected given this general distance-dependent decay. We refer to these residues with unusually strong allostery at a particular distance from the ligand as allosteric hotspots^62^. To identify allosteric hotspots in each of the four PDZ3 contexts, we first fitted an exponential decay curve to the binding energies for each construct (Fig. 6a). Here we are interested in the total change in binding energy for each mutation (ΔΔG_b_), so for the constructs with domain extension deletions we summed together the first order energies and the interaction terms. For example, for a mutation in the construct without the αC helix, the change in binding energy ΔΔG_b_(N-PDZ) = ΔΔG_b_(N-PDZ-C) + ΔΔΔG_b_(delC). For each mutation we then calculated the residual between its binding energy change and the expected binding energy change at that distance from the ligand using the fitted exponential allosteric decay function and tested if the residuals at each site were different to zero (*FDR*<0.1, one-sided t-test).

In the full-length domain (N-PDZ-C) this approach defines seven allosteric hotspot residues. Five of these sites are allosteric hotspots in all four contexts (i.e. also when each and both domain extensions are deleted; these hotspots are highlighted with red boxes in Fig. 6b; in addition, position 341 is formally a hotspot in three contexts but it also has large energy changes in PDZ-C (dashed red box in Fig. 6b,c,g).

Deletion of the N-extension does not change any of the residues defined as hotspots. However, deletion of the αC helix consistently increases the allosteric effects of mutations at five sites such that three additional sites are now defined as hotspots: R312, D357 and A375 (blue and orange boxes in Fig. 6b,c), and two hotspots (I336 and I341) are strengthened (red boxes in Fig. 6b,c). R312 is located in the first β-strand and D357 is in the fourth β-strand (cyan and orange respectively in Fig. 6d). The two residues form a salt-bridge and were previously reported as major allosteric sites in the minimal PDZ domain without both domain extensions^16^. Our results show that this strong allosteric behaviour is only observed when the αC helix extension is removed. A potential mechanism for this could be that the αC helix compensates for the energetic propagation resulting from the disruption of this salt-bridge. Position 373 located in the second α-helix was also previously reported as allosteric in the shortest domain^16^. This residue is a significant hotspot when both the N extension and αC helix are deleted and also has positive residuals in the three other contexts (yellow in Fig. 6b and Extended Data Fig. 3).

Deletion of the αC helix also results in the loss of one allosteric hotspot, with mutations in position 337 having stronger allosteric effects when the αC helix is present (dark green box in Fig. 6b). F337 directly contacts the αC helix (Fig. 6d), suggesting that mutations at this site may reduce the positive allosteric effect of αC on the binding energy. Consistent with this, mutations in F337 also only have a strong effect on the folding energy in the presence of the αC helix (Fig. 3a,b,c). The importance of this hydrophobic contact for transmitting the allosteric effect of the αC helix is also suggested by the effects of individual mutations: substituting F337 with other bulky hydrophobic amino acids (W,Y,M) does not have an allosteric effect on binding (Fig. 6c) or a strong effect on fold stability (ΔΔΔG_f_(delC) in Fig. 2e) when the αC helix is present.

In summary, the analysis of allosteric hotspots—residues where mutations have unusually strong allosteric effects at a given distance to the ligand—reveals that they are quite extensively but specifically modulated by deletion of the αC helix: two hotspots are strengthened, three additional residues become hotspots, and one hotspot is lost.

### Changes in the solvent-accessible allosteric surface

For the evolution of new regulatory mechanisms and the therapeutic targeting of proteins, solvent-accessible allosteric sites are particularly interesting^18,22^. Surface sites can be potentially engaged by physical interaction partners, including other proteins.

Only one of the allosteric hotspots identified in the full-length domain—position 341—has high solvent accessibility (relative solvent accessible surface area, rSASA = 0.239 in all four contexts). Deleting the αC helix extension results in stronger changes in binding energy for mutations in this residue (*p* = 0.000248, paired t-test comparing the average per-mutation energy of N-PDZ-C and PDZ-C (containing αC helix group) vs N-PDZ and PDZ (without αC helix group) (Fig. 6g).

Deleting the αC helix also increases the solvent exposure of an allosteric hotspot (position 329, rSASA increases from 0.141 to 0.298, Fig. 6e,f). In addition, deleting the N extension increases exposure of a residue that is only defined as an allosteric hotspot after deleting the αC deletion (position 312, rSASA increases from 0.256 to 0.375) (Fig. 6e,f).

In summary, the domain extensions also modulate the allosteric surface of the protein: removing the αC helix increases the strength of a surface-exposed allosteric hotspot and increases the accessibility of an additional hotspot. In addition, removing the N extension increases the accessibility of a residue that becomes a hotspot when the αC helix is removed. These examples illustrate the potential for domain extensions to modulate the allosteric surface of a protein, strengthening and weakening potential regulatory sites.

## Discussion

We have presented here a comprehensive quantification of how extensions to a protein domain alter its energetic and allosteric landscape. Domains are evolutionarily conserved structural units of proteins. Each instance of a domain is, however, not identical with members of a domain family quite frequently containing additional secondary structure elements or dynamic loops^6,7^. To characterise in detail the impact of domain extensions, we have used the extensively studied model system of the third PDZ domain from the human postsynaptic scaffolding protein PSD-95, and we have tested whether the energetic effects of mutations throughout the domain change when one or both extensions are removed. Measuring abundance and binding for ∼190,000 genotypes across four different domain extension contexts allowed us to quantify 6,948 energetic couplings between the domain extensions and substitutions within the domain.

The resulting comprehensive energetic and coupling landscapes provide a number of important insights. First, the effects of most mutations in most positions on stability and binding do not change when extensions are deleted. Rather, changes in the energy landscape are specific to a subset of sites. Second, sites energetically coupled to the domain extensions tend to be spatially clustered and, for couplings to the αC helix, structurally contacting and close to the helix, connecting it to the binding interface. However, strong couplings are also observed with sites distal to the domain extensions, with 7/9 sites coupled to the αC helix for folding energy and 4/6 coupled for binding energy not directly contacting it. The mechanisms underlying these distal couplings will be interesting to dissect in future work. Third, for at least two positions, domain extensions invert the sign of mutational effects, with stabilizing mutations becoming destabilizing and vice versa. Fourth, removing the domain extensions also perturbs the energy landscape of the binding interface itself, with 43% of residues (3 out of 7) in the binding interface that contribute strongly to binding being coupled to the domain extensions. Fifth, removing each domain extension also reshapes the allosteric landscape of the domain, with removal of the αC helix increasing the allosteric effects of mutations at five sites and weakening allostery at another, solvent-exposed site. Sixth, the domain extensions also alter the allosteric surface of the protein, both by altering the allosteric effects of perturbations at solvent-exposed sites and by altering the accessibility of allosteric residues.

At least for this model protein, therefore, the domain extensions profoundly and specifically alter the energetic and allosteric landscape, changing the potential for regulation and also altering evolvability.

The approach that we have taken here is quite general and can be applied to many different proteins, provided experimental selections can be performed for both function and abundance. Diverse molecular functions can be selected using growth-based assays, including protein binding^16,17^, DNA-binding^65^, and enzymatic activity^18^. As such we believe it should be possible to extend our analyses to quantify the impact of structured and dynamic extensions on the energetic and allosteric landscapes of structurally and functionally diverse proteins.

## Methods

### Media and buffers used

● LB: 10 g/L Bacto-tryptone, 5 g/L Yeast extract, 10 g/L NaCl. Autoclaved 20 min at 120°C.
● YPDA: 20 g/L glucose, 20 g/L Peptone, 10 g/L Yeast extract, 40 mg/L adenine sulphate. Autoclaved 20 min at 120°C.
● SORB: 1 M sorbitol, 100 mM LiOAc, 10 mM Tris pH 8.0, 1 mM EDTA. Filter sterilized (0.2 mm Nylon membrane, ThermoScientific).
● Plate mixture: 40% PEG3350, 100 mM LiOAc, 10 mM Tris-HCl pH 8.0, 1 mM EDTA pH 8.0. Filter sterilised.
● Recovery medium: YPD (20 g/L glucose, 20 g/L Peptone, 10 g/L Yeast extract) + 0.5 M sorbitol. Filter sterilised.
● SC –URA: 6.7 g/L Yeast Nitrogen base without amino acid, 20 g/L glucose, 0.77 g/L complete supplement mixture drop-out without uracil. Filter sterilised.
● SC –URA/MET/ADE: 6.7 g/L Yeast Nitrogen base without amino acid, 20 g/L glucose, 0.74 g/L complete supplement mixture drop-out without uracil, adenine and methionine. Filter sterilised.
● Competition medium: SC –URA/MET/ADE + 200 ug/mL methotrexate (MERCK LIFE SCIENCE), 2% DMSO.
● DNA extraction buffer: 2% Triton-X, 1% SDS, 100mM NaCl, 10mM Tris-HCl pH8, 1mM EDTA pH8.

### Library design

In each context we constructed a library of single and double aa variants spanning 66 residues of the canonical PDZ domain (residues 312 to 377). From this region, we selected double mutants based on several criteria to enrich for residues likely to be energetically coupled. First we included all non-adjacent interacting residue pairs within 5Å of each other in either the bound (PDB ID = 1BE9) or the unbound (PDB ID = 1BFE) structures^41^, excluding consecutive residues in the linear amino acid sequence (a total of 100 pairs). Second, we added all second shell contacts, defined as residues within 5Å of any binding interface residue (5 pairs), as well as adjacent pairs in which only one of the residues is part of the binding interface.

To capture known allosteric connectivity, we also incorporated all residue pairs within the previously characterized wire-like connected set of allosteric residues outside of the binding interface (residues R312, I336, and D357)^16^ (2 pairs). Additionally, to explore long-range allosteric communication, we also included all residue pairs including a distal allosteric residue (R312 and D357) and any binding interface residue with a weighted mean absolute ΔΔG above the average for binding interface residues (12 pairs).

Finally, we included further 10 randomly selected non-adjacent pairs of residues separated by more than 5Å of distance, and 10 randomly selected pairs of adjacent residues at less than 5Å.

Each pair of sites was co-mutated using NNK codons at both positions, generating a combinatorial of all 19 amino acid substitutions at each site. Each residue pair was encoded as a separate oligonucleotide, and all oligonucleotides were purchased as two pools from IDT (Supplementary Table 4) one to be cloned into domains containing N terminal (N-PDZ and N-PDZ-C, with homology to Nterm (5’-GACATTCCCCGAGAACCG-3’)) and another to be cloned into domains without it (PDZ-C and PDZ, with homology to the linker (5’-GGAGGTGGAGCTAGCCCG-3’)).

### Plasmid construction

The wild type-containing plasmids, pGJJ210 (PDZ-C), and pGJJ211 (N-PDZ-C) were built by Gibson reaction (in house preparation) from plasmid pGJJ111 (PDZ) described in^16^. pGJJ527 (N-PDZ) plasmid was built by PCR amplification and blunt ligation (T4 ligase (NEB)) from pGJJ211 (N-PDZ-C). Backbone bPCA (pGJJ518) plasmid is described in^35^ and aPCA (pGJJ045) plasmid are described in^16^. All plasmids and primers used in the study are described in (Supplementary Table 1 and Supplementary Table 2).

### Library construction

Libraries were constructed in two steps. First, the oligonucleotide library purchased from IDT was assembled by Gibson into the mutagenesis plasmids already containing the wild-type version of each domain pGJJ111 (PDZ), pGJJ210 (PDZ-C), pGJJ211 (N-PDZ-C) and pGJJ527 (N-PDZ). Libraries were then cloned into the yeast plasmids aPCA (pGJJ045) and bPCA (pGJJ518) by digestion/ligation.

For the first step the library amplification was performed in 12 PCR cycles (Supplementary Table 2) with Q5 polymerase (New England Biolabs). dsDNA product was then incubated with ExoSAP-IT (Applied Biosystem) to get rid of the remaining ssDNA and purified with MinElute columns (Qiagen). The plasmid mutagenesis backbones (Supplementary Table 1) were also linearised (Supplementary Table 2), incubated with DpnI to remove the template and gel purified using QIAquick gel extraction kit (Qiagen). 100ng of backbone in a molar ratio 1:5 with the insert was incubated for 5h at 50°C with a gibson mix 2x prepared in house. The reaction was desalted by dialysis with membrane filters (MF-Millipore) for 1h and concentrated 4X using SpeedVac concentrator (Thermo Scientific). DNA was then transformed into NEB 10β High-efficiency Electrocompetent *E. Coli*. Cells were allowed to recover in SOC medium (NEB 10β Stable Outgrowth Medium) for 30 minutes and later transferred to LB medium with ampicillin overnight. A fraction of cells was also plated into ampicillin+LB+agar plates to estimate the total number of transformants. 100 mL of each saturated *E. coli* culture were harvested the next morning to extract the mutagenesis plasmid library using the Qiagen Plasmid Plus Midi Kit (QIAGEN). To transfer the final libraries into the assay plasmids, the libraries inside each mutagenesis plasmid were digested with NheI and HindIII, and later gel purified (MinElute Gel Extraction Kit, QIAGEN). Clean libraries were then ligated into pGJJ045 or pGJJ518+CRIPT digested plasmids using T4 ligase (New England Biolabs) with temperature-cycle ligation, following manufacturer instructions. The ligation was desalted by dialysis using membrane filters for 1h, concentrated 4X using a SpeedVac concentrator (Thermo Scientific) and transformed into NEB 10β High-efficiency Electrocompetent *E. coli* cells. 50mL of grown culture was collected the next morning for DNA extraction using the Qiagen Plasmid Plus Midi Kit (QIAGEN).

### Methotrexate yeast selections

The yeast selection assay was described in^16^. Three independent pre-cultures of BY4742 were grown in 50 mL standard YPDA at 30°C overnight. The next morning, the cultures were diluted into 400 mL of pre-warmed YPDA at an OD_600nm_ = 0.3 and incubated at 30°C for 4 hours. Cells were then harvested and centrifuged for 5 minutes at 3,000g, washed with sterile water and SORB medium, resuspended in 17.2 mL of SORB and incubated at room temperature for 30 minutes. After incubation, 350 μL of 10mg/mL boiled salmon sperm DNA (Agilent Genomics) and 7 μg of plasmid library were added to each tube of cells and mixed gently. Two tubes with 35 mL of Plate Mixture were prepared for each replicate. The same amount of cells in SORB were added to each tube and were incubated at room temperature for an additional 30 minutes.

3.5 mL of DMSO was added to each tube and the cells were then heat shocked at 42°C for 20 minutes (inverting tubes from time to time to ensure homogenous heat transfer). After heat shock, cells were centrifuged and re-suspended in ∼50 mL of recovery media and allowed to recover for 1 hour at 30°C. Cells were then centrifuged, washed with SC-URA medium and re-suspended in 400mL SC –URA. 10 uL were plated on SC –URA Petri dishes and incubated for ∼48 hours at 30°C to measure the transformation efficiency. The independent liquid cultures were grown at 30°C for ∼48 hours until saturation. Saturated cells were diluted again to OD_600nm_ = 0.05 in SC –URA/MET/ADE media and allowed to grow 5 generations until OD_600nm_ = 1.6 at 30°C and 200rpm. A fraction of the culture was then used to inoculate 400mL of competition media containing methotrexate at a starting OD_600nm_ = 0.05, and the rest was harvested and pellets frozen and stored as INPUT. Cells in competition media were allowed to grow for 5 generations, collected and frozen and stored as OUTPUT.

### DNA extractions and plasmid quantification

The DNA extraction protocol used was described in^16^. Cell pellets (one for each experiment input/output replicate) were re-suspended in 2 mL of DNA extraction buffer, frozen by dry ice-ethanol bath and incubated at 62°C water bath twice. Subsequently, 2 mL of Phenol/Chloro/Isoamyl 25:24:1 (equilibrated in 10mM Tris-HCl, 1mM EDTA, pH8) was added, together with 1g of acid-washed glass beads (Sigma Aldrich) and the samples were vortexed for 10 minutes. Samples were centrifuged at RT for 30 minutes at 4,000 rpm and the aqueous phase was transferred into new tubes. The same step was repeated twice. 0.2 mL of NaOAc 3M and 4.4 mL of pre-chilled absolute ethanol were added to the aqueous phase. The samples were gently mixed and incubated at –20°C at least for 30 minutes. After that, they were centrifuged for 30 min at full speed at 4°C to precipitate the DNA. The ethanol was removed and the DNA pellet was allowed to dry overnight at RT. DNA pellets were resuspended in 0.6 mL TE 1X and treated with 5 uL of RNaseA (10mg/mL, Thermo Scientific) for 30 minutes at 37°C. To desalt and concentrate the DNA solutions, QIAEX II Gel Extraction Kit was used (50 µL of QIAEX II beads, QIAGEN). The samples were washed twice with PE buffer and eluted twice by 125 µL of 10 mM Tris-HCI buffer, pH 8.5. Finally, plasmid concentrations in the total DNA extract (that also contained yeast genomic DNA) were quantified by qPCR using the primer pair oGJJ152-oGJJ153, that binds to the ori region of the plasmids.

### Sequencing library preparation

The library preparation protocols for illumina sequencing was previously described in^16^. Briefly, the sequencing libraries were constructed in two consecutive PCR reactions. The first PCR (PCR1) was designed to amplify the mutated protein of interest and to increase the nucleotide complexity of the first sequenced bases by introducing frame-shift bases between the adapters and the sequencing region of interest. Two different forward pools were used depending on the presence (oGJJ800 pool) or absence (oGJJ052 pool) of N terminal extension in the domain (Supplementary Table 2). The second PCR (PCR2) was necessary to add the remainder of the Illumina adapter and demultiplexing indexes (dual indexing (or just reverse index for PDZ_aPCA, Supplementary Table 3) unique to each sample. All samples were pooled in an equimolar ratio and gel purified using the QIAEX II Gel Extraction Kit. The amplicons were subjected to Illumina paired end 2×150 sequencing on a NextSeq2000 (or NextSeq500 for PDZ_aPCA library) instrument at the CRG Genomics facility.

### Sequencing data processing

Raw FastQ files were processed using DiMSum pipeline^66^ (https://github.com/lehner-lab/DiMSum). FASTQ files corresponding to the same sample but originating from different sequencing runs or lanes were treated as technical replicates.

Two rounds of DiMSum runs were performed. In the first round, all input and output samples were analyzed together with wild-type sequencing data (pGJJ068 or pGJJ518 as template, depending on the absence or presence of N extension respectively). The FastQ file for the wild type sample was processed identically to those of the replicate Input/Output samples with permissive base quality thresholds (“vsearchMinQual = 5” and “vsearchMaxee = 1000”). Read counts for all variants were then adjusted by subtracting the expected number of sequencing errors derived from the wild-type-only sample and proportional to the total sequencing library size of each sample. Once the new variant count files were calculated, the next dimsum run was performed. Restarting at stage 4, the modified variant count files were given as input (option “countPath”). Together with this, barcode-variant identity files were also provided to restrict the analysis to designed variants only (option “barcodeIdentityPath”). Filtering thresholds were informed by the input count distribution plots (Stage 4.1) and expected read counts of variants arising from sequencing errors, included in the DiMSum report^66^. The Dimsum output “…fitness_replicates.RData” files containing the fitness and fitness error estimates were used for further data analysis.

### Thermodynamic modeling with MoCHI

We used MoCHIv0.9^31^ (https://github.com/lehner-lab/MoCHI) to fit a thermodynamic model using the corresponding 8 aPCA and bPCA datasets (2 molecular phenotypes x 4 domain boundaries) simultaneously.

The software is based on our previously described genotype-phenotype modeling approach^31^ with additional functionality and improvements for ease-of-use and flexibility.

We model protein folding as an equilibrium between two states: unfolded and folded, and protein binding as an equilibrium between three states: unfolded and unbound, folded and unbound, and folded and bound. We assume that the probability of the unfolded and bound state is negligible. This model fits single mutation Gibbs free energy of folding (ΔG_f_) and binding (ΔG_b_) effects, and works under the assumption that free energy changes of folding (ΔΔG_f_) and binding (ΔΔG_b_) are additive. This means that the effect of mutations from a multi-mutant are simply the sum of effects of the single mutations.

MoCHI parameters were set to default as explained in previous work^16–18,31^ to specify a neural network architecture consisting of additive trait layers (free energies) for each biophysical trait to be inferred (folding or folding and binding for aPCA or bPCA, respectively), as well as one linear transformation layer per observed phenotype. The specified non-linear transformations “TwoStateFractionFolded” and “ThreeStateFractionBound” derived from the Boltzmann distribution function relate energies to proportions of folded and bound molecules respectively (see Fig. 2a and Fig. 2b). The inclusion of first-, second– or third-order (energetic couplings) model coefficients in the models was specified using the “max_interaction_order” option.

As input for MoCHI we used raw fitness scores from both single and double mutants. DiMSum output tables were manually adjusted so that we dummy-encoded the domain extension deletions as two additional amino acids appended to the canonical domain sequence (the target of amino acid substitution mutagenesis).

The design of our library, comprising 144 mutation pairs across 61 amino acids, in 4 different extension contexts, and two assays, yields a theoretical total of more than 110000 possible model terms to be fitted simultaneously, including 103,968 second-order terms corresponding to couplings between amino acid substitutions (ΔΔΔG(A,B)) for binding and folding energies. Due to memory constraints, modeling all these features simultaneously is not feasible. Therefore, we used a three-round procedure to iteratively reduce the number of energy terms fitted, removing substitution coupling terms with values close to zero. This resulted in a final list of 8895 coefficient features per assay (Extended Data Fig. 2c). All the.txt files are available at: https://doi.org/10.5281/zenodo.16411462.

Below and in Extended Data Fig. 2c are explained the details for the features filtering:

### Round 1

This round fits 24x 1st and 2nd order MoCHI models to identify substitution coupling features with values close to 0. For each model fitted, the starting features list is included in the file features_all_55043.txt. This initial list includes, per assay:

– 1 x WT (ΔG_wt_)
– 1158 x first order s (s = aminoacid substitutions) (ΔΔG)
– 1 x first order delN (deletion of extra N terminal) (ΔΔG(delN))
– 1 x first order delC (deletion of extra C terminal) (ΔΔG(delC))
– 1158 x second order s-delN interaction (ΔΔΔG(delN))
– 1158 x second order s-delC interaction (ΔΔΔG(delC))
– 1 x second order delN-delC interaction (ΔΔΔG(delNC))
– 51566 second order s-s interactions (ΔΔΔG(A,B))

Each MoCHI is ran with the following parameters:

–– features=features_all_55043.txt
–– model_design=PDZ_model_design_bothshared_interactions.txt
–– max_interaction_order 2
–– l1_regularization_factor “0.01,0.001,0.0001”
–– downsample_interactions 2500

The final parameter is used to downsample the 2nd order features fitted in each MoCHI run to a random subset of 2,500, but keeping all 1st order terms. As a result, the first-round MoCHI runs produce 24 coefficient weight tables.

These tables are then merged, and the weights calculated across multiple MoCHI models are aggregated as a weighted mean and associated error for each energy term. This results in the following two tables:

– merge_table_s_s_Folding_from_55043.txt
– merge_table_s_s_Binding_from_55043.txt

For each of these two tables separately, we identify A-B couplings whose absolute ΔΔΔG(A,B) values are smaller than half the standard deviation of the ΔΔΔG(A,B) dataset, as these are considered to be ∼0: |ΔΔΔG(A,B)|< std_dev * 0.5.

From the original list of features (features_all_55043.txt), we remove all A-B couplings that were classified as ∼0 in both assays. The result is a reduced feature set saved in: features_2nd_round_24622.txt.

### Round 2

The same procedure is then repeated in a second round of 30 MoCHI fits, using as input the filtered list of 1st and 2nd order terms from round 1 (features_2nd_round_24622.txt).

Each MoCHI is ran with the following parameters:

–– features=features_2nd_round_24622.txt
–– model_design=PDZ_model_design_bothshared_interactions.txt
–– max_interaction_order 2
–– l1_regularization_factor “0.01,0.001,0.0001”
–– downsample_interactions 2500

As in round1, in each model, 2nd order features are randomly subsampled to 2,500, but keeping all 1st order terms. As a result, these second-round MoCHI runs produce 30 coefficient weight tables.

These tables are then merged, and the weights calculated across multiple MoCHI models are aggregated as a weighted mean and associated error for each energy term. This results in the following two tables:

– merge_table_s_s_Folding_from_24622.txt
– merge_table_s_s_Binding_from_24622.txt

Again, for each of these two tables separately, we identify A-B couplings whose absolute ΔΔΔG(A,B) values are smaller than half the standard deviation of the ΔΔΔG(A,B) dataset, as these are considered to be ∼0: |ΔΔΔG(A,B)|< std_dev * 0.5.

From the round 1 output list of features (features_2nd_round_24622.txt), we remove all A-B couplings that were classified as ∼0 in both assays. The result is a reduced feature set saved in: features_3rd_round_9460.txt.

### Round 3

The same procedure is then repeated in a third round of 30 MoCHI fits, using as input the filtered list of 1st and 2nd order terms from round 2 (features_3rd_round_9460.txt).

Each MoCHI is ran with the following parameters:

–– features=features_3rd_round_9460.txt
–– model_design=PDZ_model_design_bothshared_interactions.txt
–– max_interaction_order 2
–– l1_regularization_factor “0.01,0.001,0.0001”
–– downsample_interactions 2500

These tables are then merged, and the weights calculated across multiple MoCHI models are aggregated as a weighted mean and associated error for each energy term. This results in the following two tables:

– merge_table_s_s_Folding_from_9460.txt
– merge_table_s_s_Binding_from_9460.txt

Again, for each of these two tables separately, we identify A-B couplings whose absolute ΔΔΔG(A,B) values are smaller than half the standard deviation of the ΔΔΔG(A,B) dataset, as these are considered to be ∼0: |ΔΔΔG(A,B)|< std_dev * 0.5.

From the round 2 output list of features (features_3rd_round_9460.txt.), we remove all A-B couplings that are not found after the third round in both assays. The result is a reduced feature set saved in: features_3rd_round_9460_final.txt.

### Final MoCHI models

After all A-B couplings ∼0 filtered, we add to the final reduced feature set the third order interactions for aa substitutions and removal of both N and C (ΔΔΔΔG(delNC)). This final features list is saved in third_order_features_s_s.txt, which includes:

– 1 x WT (ΔG_wt_)
– 1158 x first order s (s = aminoacid substitutions) (ΔΔG)
– 1 x first order delN (deletion of extra N terminal) (ΔΔG(delN))
– 1 x first order delC (deletion of extra C terminal) (ΔΔG(delC))
– 1158 x second order s-delN interaction (ΔΔΔG(delN))
– 1158 x second order s-delC interaction (ΔΔΔG(delC))
– 1 x second order delN-delC interaction (ΔΔΔG(delNC))
– 4260 second order s-s interactions (ΔΔΔG(A,B))
– 1158 x third order s-delN-delC interactions (ΔΔΔΔG(delNC))

Each 30x final MoCHI run includes the following parameters:

–– features=third_order_features_s_s.txt
–– model_design=PDZ_model_design_bothshared_interactions.txt
–– max_interaction_order 3
–– l1_regularization_factor “0.01,0.001,0.0001”
–– downsample_interactions 1500,1000

In this case fitted terms are downsampled to a random subset of 1500 for 2nd order terms, and 1000 for 3rd order terms. As a result, these final-round MoCHI runs produce 30 coefficient weight tables.

These tables are finally merged, and the weights calculated across multiple MoCHI models are aggregated as a weighted mean and associated error for each energy term. This yields the final dataset of folding and binding energetic effects for 1st, 2nd, and 3rd order mutation interactions used in this study.

### Identification of mutations with significant changes in energy terms and enrichments by residue

The energy values obtained from MoCHI fits were used for statistical testing to identify mutations with significant changes (|ΔΔG|, |ΔΔΔG| or |ΔΔΔΔG|) > 0) in folding or binding. We performed Z-test and p-values were corrected for multiple testing using the false discovery rate (FDR) method Benjamini-Hochberg, with a significance threshold of *FDR* < 0.05. We applied an additional cut-off to avoid false positives of |energy-term|_mutation_ > median |energy-term| _dataset_ + Median Absolute Deviation(MAD).

Enrichments of mutations identified as significant in individual residues were tested using Fisher’s exact test (*FDR* < 0.1) compared to the rest of the residues.

### Protein structure metrics

All the calculations were done using the PDB:1BE9 structure. Minimum ligand-residue distances were calculated as the shortest distance between any side chain heavy atom of the residue and the ligand side chain heavy atom. Relative solvent accessible surface area (rSASA) was calculated for modified versions of PDB:1BE9 resulting from trimming residues 303 to 402 (N-PDZ-C), 311 to 402 (PDZ-C), 303 to 394 (N-PDZ) or 311 to 394 (PDZ) using freeSASA python package (v2.2.1)^67^. All residues with rSASA>0.2 were defined as solvent exposed.

## Structure visualizations

ChimeraX^68^ was used to visualise all protein structures presented. Contacts displayed in these structures are drawn using the contacts command with default settings.

## Allosteric hotspots

To quantify the dependence of mutation effects on the distance from the ligand, we computed the minimum heavy atom side chain distance between each residue and the CRIPT ligand from the PDB:1BE9 structure. To quantify this dependence, an exponential decay function was fitted to the data, with initial parameter estimates obtained after transformation to a linear function (*y* = *a* · *e^bx^*to linear log(y) = log(a)+b⋅x in order to obtain the a and b coefficients). In exponential decay model *y = a · e^bx^*, *a* represents the estimated |ΔΔG_b_| at distance 0 Å from the ligand, *b* is the decay rate and x is the minimum heavy atom side chain distance to the ligand. The model was fit to all data points excluding those corresponding to binding interface residues (binding interfaces were defined as any residue within 5 Å of the ligand).

To identify allosteric hotspots, we first calculated the expected ΔΔG_b_ at each residue based on its minimum side-chain heavy atom distance to the ligand, using the previously fitted exponential decay model. Residuals were then computed as the difference between observed and expected ΔΔG_b_ values. Residues were classified as allosteric hotspots if their residual distribution was significantly greater than zero, using a one-sided t-test with FDR correction (*FDR* < 0.1). Sometimes residuals of mutations that are far away from the ligand are different to zero due to the low value of the curve for far away residues (example of mutations of residue R312 in pink in Fig. 6a). To avoid false positives do to this we set up a cut off of energy values for mutations |ΔΔG_b_| > median ΔΔG_b_ + MAD (dashed blue line in Fig. 6a) and took into account only residues enriched for mutations above that cut off as explained before. This is the same result as applying the residuals t-test only to residues enriched for allosteric mutations in previous sections. For example, of the eight residues outside the binding interface shown before for the full-length PDZ domain (FDR<0.1, FET, Fig. 5b) only one (373) does not appear as an allosteric hotspot now (Extended Data Fig. 3 where in pink are the mutations for residue 373). Mutations are above the cut off in all the 4 domains (FDR<0.05, Z-test and |ΔΔG_b_|>(median ΔΔG_b_ + Median Absolute Deviation(MAD)) but only for the shorter PDZ they are also significantly above the decay curve (*FDR*<0.1, one-sided t-test).

## Figure legends

**Extended Data Figure 1:**
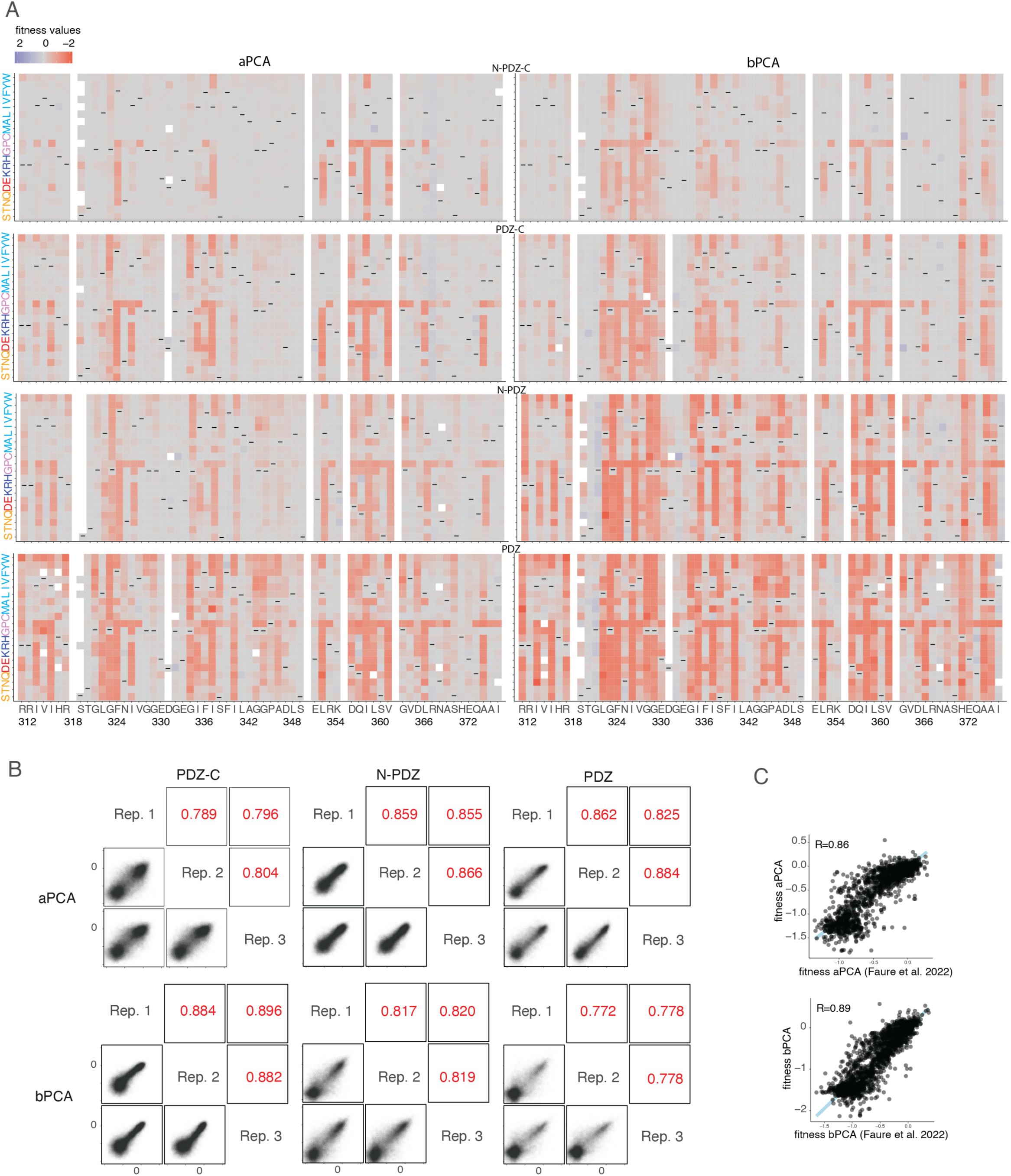
Binding and Abundance selections. **a.** Heatmaps for fitness effects of single amino acid substitutions from aPCA (left) and bPCA (right). Dashes indicate wild-type residues. **b.** Scatter plots showing the reproducibility of fitness estimates from aPCA (top) and bPCA (bottom). Pearson’s R indicated in red. Rep., replicate. **c.** Correlation of fitness values with previous measurements^16^ for the canonical domain without domain extensions for aPCA (top) and bPCA (bottom), respectively, Pearson’s R indicated.

**Extended Data Figure 2:**
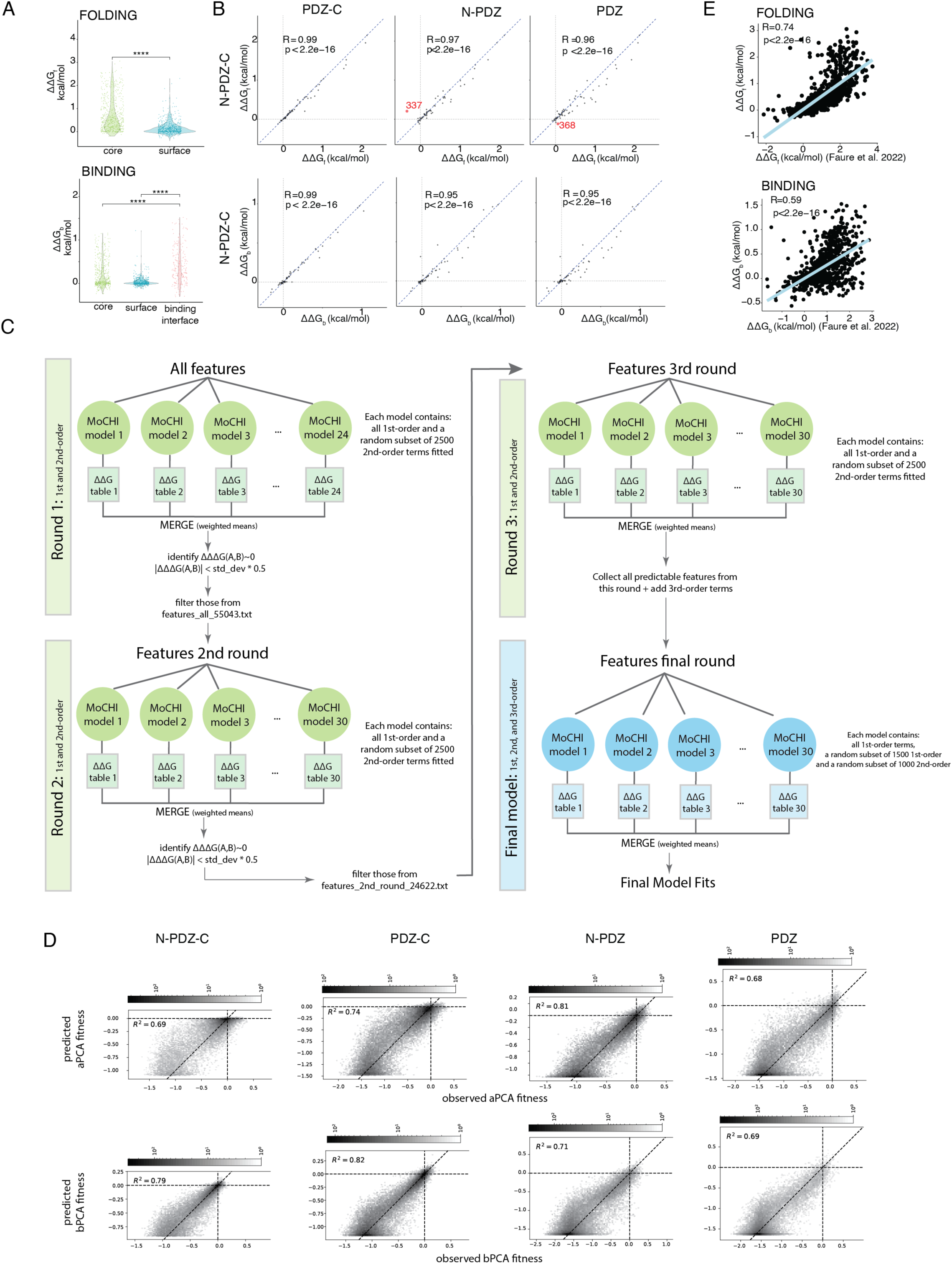
Energy landscapes. **a.** Violin plots showing ΔΔG_b_ for mutations in core and surface for Folding (left) and in core, surface and binding interface for Binding (right). (surface defined by rSASA > 0.2 in full length domain and binding interface defined as residues with minimum side chain heavy atom distance to the ligand less than 5 Å). **** means *p* < 0.0001(t-test). **b.** Correlation of free energy changes (ΔΔG) between full-length N-PDZ-C and domains with extensions deleted for folding (ΔΔG_f_, panels on the top) or binding (ΔΔG_b_, panels on the bottom). In red are labeled residues that change the sign of the energy compared to full-length (337 from positive to negative in N-PDZ and 368 from negative to positive in PDZ-C). R = Pearson’s coefficient. **c.** Diagram explaining the process of filtering features to fit in MoCHI. **d.** Example of MoCHI performance. Plots for each assay (aPCA: top, bPCA: bottom) and PDZ domain. Shown is one example of the 30 x MoCHI runs since the variance explained is always the same for the 30 runs.

**Extended Data Figure 3:**
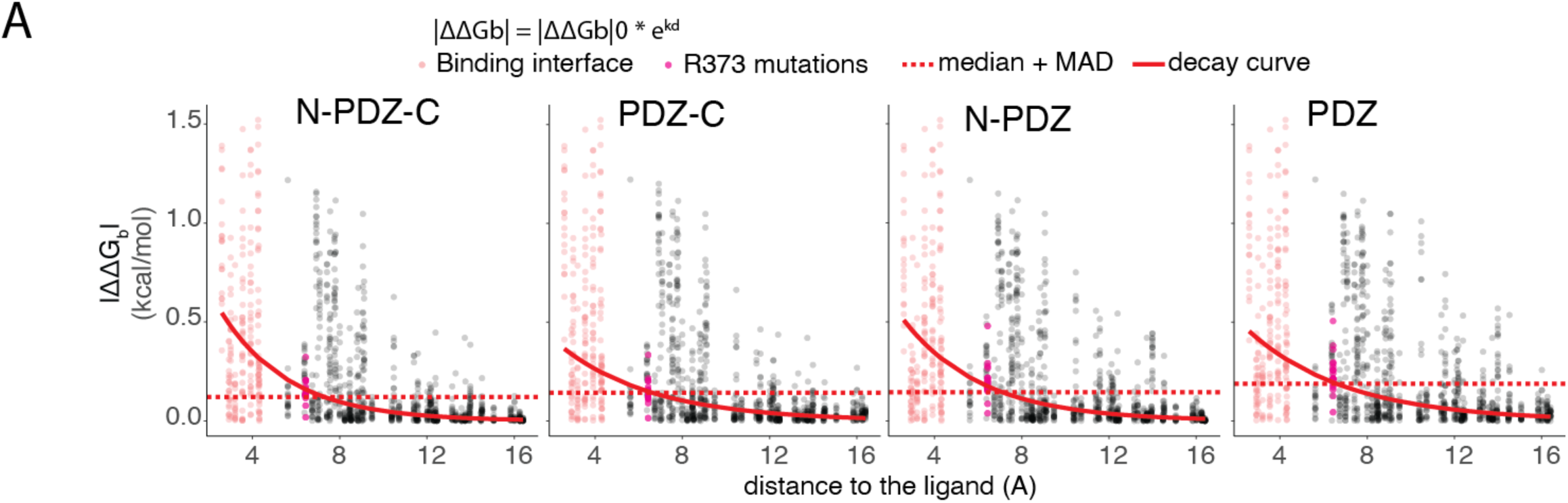
Allostery. **a.** Exponential decay fits to the relationship between |ΔΔG_b_| to the minimum heavy atom distance to the ligand. |ΔΔG_b_|_0_=starting ΔΔG_b_ at distance d = 0, k = decay rate, d = distance from the ligand. Points are red if in the binding interface (minimal side chain heavy atom distance to ligand < 5 Å). Magenta points correspond to mutations of the residue 373. Dash blue line is in |ΔΔG_b_| = median + MAD, see Materials and Methods.

## Acknowledgements

This work was funded by a European Research Council (ERC) Advanced Grant (883742) under the European Union’s Horizon 2020 research and innovation programme, the Spanish Ministry of Science and Innovation (PID2020-118723GB-I00), Wellcome (220540/Z/20/A), Departament de Recerca i Universitats de la Generalitat de Catalunya (AGAUR, 2021 SGR 01226). A.J.F. was funded by a Ramón y Cajal fellowship (RYC2021-033375-I) financed by the Spanish Ministry of Science and Innovation (MCIN/AEI/10.13039/501100011033) and the European Union (NextGenerationEU/PRTR). A.M.A. was funded in part by a fellowship from “laCaixa” Foundation (ID 100010434, fellowship code B006052). T.Z. was funded by a EMBO Long-term Postdoctoral Fellowship (ALTF 525-2021) and a Marie Skłodowska-Curie Postdoctoral Fellowship (European Union’s Horizon Europe under the grant, GPIDR, 101068134). We acknowledge support of the Spanish Ministry of Science and Innovation through the Centro de Excelencia Severo Ochoa (CEX2020-001049-S, MCIN/AEI /10.13039/501100011033), and the Generalitat de Catalunya through the CERCA programme. We are grateful to the CRG Core Technologies Programme for their support and assistance in this work. We acknowledge all members of the Lehner lab for feedback during the project, particularly Toni Beltran and Albert Escobedo for advice with data analysis.

## Data availability

All DNA sequencing data have been deposited in the Gene Expression Omnibus (GEO) with accession number GSE299757. Files to reproduce all figures in this work are available at: https://doi.org/10.5281/zenodo.16411462

## Code availability

Source code to reproduce the analyses is available at https://github.com/lehner-lab/pdzext

## Author contributions

C.H.-C. performed all experiments and analyses and made all the figures; C.H.-C. and A.J.F. developed the model fitting approach; A. M.-A. designed the library; T.Z. performed pilot experiments. B.L. conceived and supervised the project. C.H.-C. and B.L. wrote the manuscript with input from all authors.

